# Polydopamine Nanoparticles as Mimicking RPE Melanin for the Protection of Retinal Cells Against Blue Light-Induced Phototoxicity

**DOI:** 10.1101/2024.01.15.575719

**Authors:** Yong-Su Kwon, Min Zheng, Alex I. Smirnov, Zongchao Han

## Abstract

Exposure of the eyes to blue light can induce the overproduction of reactive oxygen species (ROS) in the retina and retinal pigment epithelium (RPE) cells, potentially leading to pathological damage of age-related macular degeneration (AMD). While the melanin in RPE cells absorbs blue light and prevents ROS accumulation, the loss and dysfunction of RPE melanin due to age-related changes may contribute to photooxidation toxicity. Herein, we present a novel approach utilizing a polydopamine-replenishing strategy via a single-dose intravitreal (IVT) injection to protect retinal cells against blue light-induced phototoxicity. To investigate the effects of overexposure to blue light on retinal cells, we created a blue light exposure Nrf2-deficient mouse model, which are susceptible to light-induced retinal lesions. After blue light irradiation, we observed retina degeneration and an overproduction of ROS. The Polydopamine-replenishing strategy demonstrated effectiveness in maintaining retinal structural integrity and preventing retina degeneration by reducing ROS production in retinal cells against the phototoxicity of blue light exposure. Our findings highlight the potential of polydopamine as a simple and effective replenishment for providing photoprotection against high-energy blue light exposure.

**Graphical Abstract:** 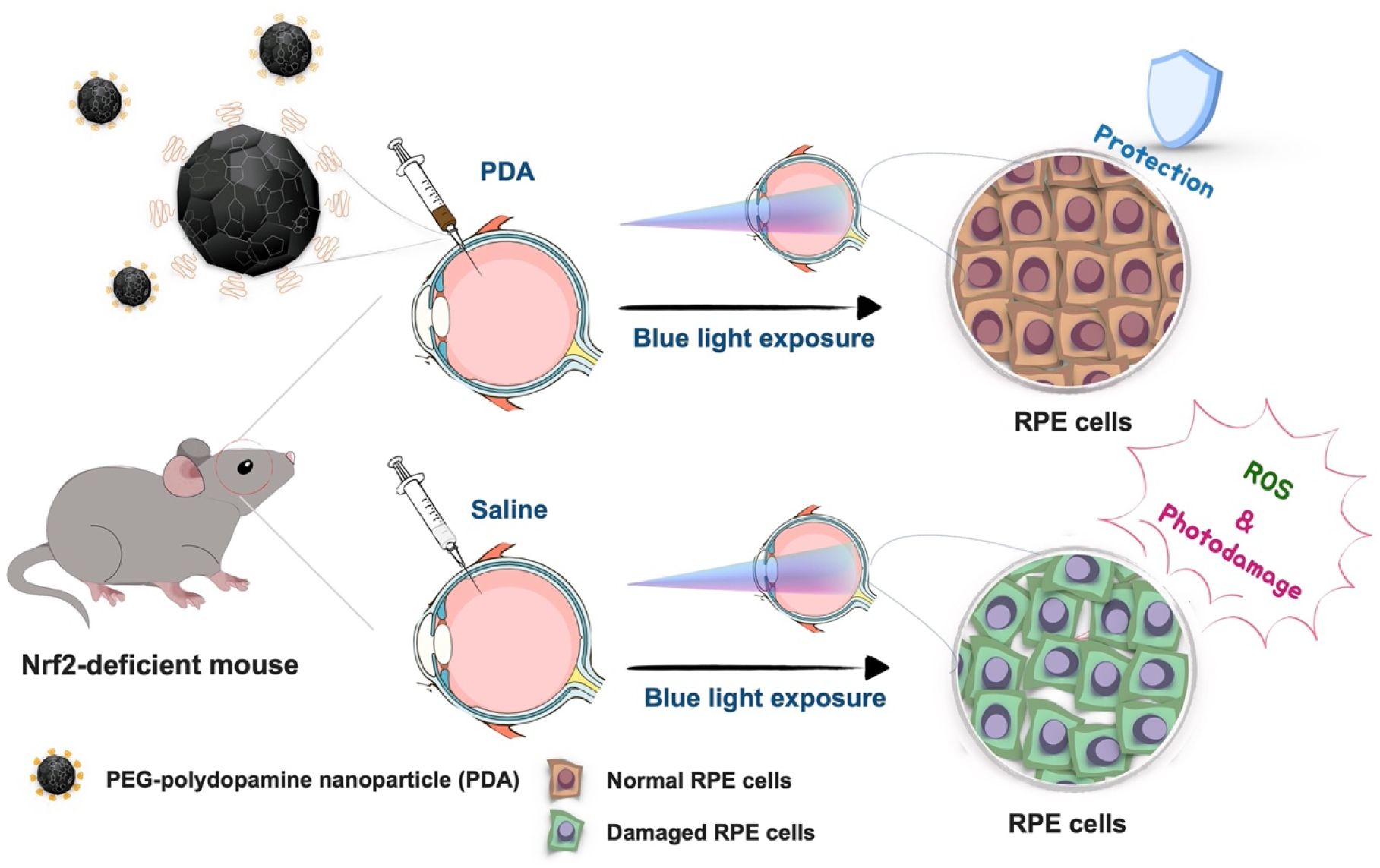

The graphic illustration of PDA-mediated photoprotection strategies to mimic natural melanin in RPE cells against blue light-induced phototoxicity in an Nrf2-deficient mouse model via a single-dose intravitreal (IVT) administration.

## 1. Introduction

Age-related macular degeneration (AMD) is a significant contributor to central vision loss, primarily affecting the elderly population and often leading to severe vision impairment or even legal blindness. With a far-reaching impact, AMD is projected to affect approximately 288 million individuals worldwide by 2040[1]. The exact cause of AMD remains not fully understood, but it is considered to be linked to oxidative stress and the depletion of systemic antioxidant capacity[2,3].

The widespread use of modern technological devices, particularly commercially available light-emitting diodes (LEDs) with intense emission, has led to a significant increase in blue light exposure and raised growing concerns about the potential implications. The light undergoes a sequential filtering process as it traverses each layer of intraocular structures, including the cornea, lens, and vitreous humor[4]. These advantageous absorption characteristics limit the potentially damaging ultraviolet radiation to retinal pigment epithelium (RPE) cells. However, the range of blue light (between 415 nm and 455) is distinguished by its strong ability to reach the RPE layer, in contrast to other visible light components, with relatively little attenuation[5,6].

Lipofuscin is a metabolic product of phagocytosis in the RPE cells of photoreceptor outer segments, which includes N-retinylidene-N-retinylethanolamine (A2E) with autofluorescence and phototoxicity[7,8]. A2E increases with age and is not easily degraded, leading to the abnormal accumulation of lipofuscin observed in dry AMD[9]. In particular, exposure of A2E to blue light could induce the production of substantial reactive oxygen species (ROS) such as superoxide anions, singlet oxygen, and hydrogen peroxide due to structural changes to highly reactive A2E-epoxides form[10]. Consequently, this may induce RPE cells apoptosis or result phototoxicity in retinal cells, thus emphasizing the critical importance of robust antioxidant defenses in maintaining retinal health[11]. Furthermore, the retina, one of the most oxygen-consuming tissues in the human body, possesses an abundance of mitochondria, which could potentially cause oxidative stress through blue light stimulation due to the presence of blue light-absorbing photosensitizers in mitochondria, such as flavin and porphyrins[12–14].

Melanin is a natural pigment that is present in many tissues such as RPE cells. RPE melanin serves a photoprotective role, contributing to protecting the light-sensitive ocular tissues against oxidative stress and phototoxicity[15,16]. However, the quality and quantity of RPE melanin decline with increasing age[17]. These factors could lead to an increased risk of excess production of ROS in retinal cells, contributing to the pathogenesis of AMD. Polydopamine nanoparticles, which share similar physical and chemical properties with natural eumelanin in RPE cells[18–20], possess excellent biocompatibility and may provide a new opportunity for supporting the performance improvement of natural RPE melanin.

In this work, we explored the efficacy of PDA with antioxidative and photoprotective properties as a substitute for natural melanin in protecting retinal cells against photodamage induced by LEDs blue light in an Nrf2-deficient mouse model. Our findings described in the present study provide new insights into the use of PDA with biomimetic photoprotective characteristics for replenishing natural RPE melanin.

## 2. Results

### 2.1 Synthesis and Characterization of polydopamine nanoparticles

The various steps involved in the synthesis of PEGylated-polydopamine nanoparticles (PDA) through the dopamine polymerization process are outlined in **Figure 1a**. In brief, dopamine monomers underwent auto-oxidation at pH 8.5 in the presence of hydroxide ions, resulting in the formation of spherical bare-PDA. Subsequently, the PEGylated shell on bare-PDA was formed using polyethylene glycol–thiol (mPEG–SH) via Michael addition reactions[21]. The morphology of bare-PDA revealed uniform spherical shapes with an average size of approximately 97 ± 6 nm **(Figure 1b and Figure S1a in the Supporting Information)** and showed no morphological changes in PDA after modification with PEG-SH on the surface, as confirmed by TEM (**Figure 1c and Figure S1b**). The hydrated size of bare-PDA was 122.4 ± 4 nm, while PDA measured 131.32 ± 3.3 nm due to surface modification with PEG **(Figure 1d)**. The zeta potential of bare-PDA exhibited – 22.4 mV, which changed to – 7.8 mV after PEGylation as shown in **Figure 1e**, indicating the successful preparation of PDA. The colloidal stability results indicated that bare-PDA was precipitated in pig vitreous within 14 days **(Figure S2 supporting information)**, whereas PDA was significantly enhanced colloidal stability through surface modification with PEG, demonstrating that PDA maintains excellent dispersity in the pig vitreous for 2 months **(Figure 1f)**. As shown in **Figure 1g**, we further confirmed that PDA dispersions in pig vitreous showed similar absorption spectra compared with those of the aqueous solution, indicating the well-preservation of PDA molecules in pig vitreous solution. The absorbance at 808 nm exhibited a good linear relationship with different concentrations of PDA **(Figure S3 supporting information)**, providing a foundation for accurately quantifying PDA concentration for further studies **(Figure 1h)**.

**Figure 1.**
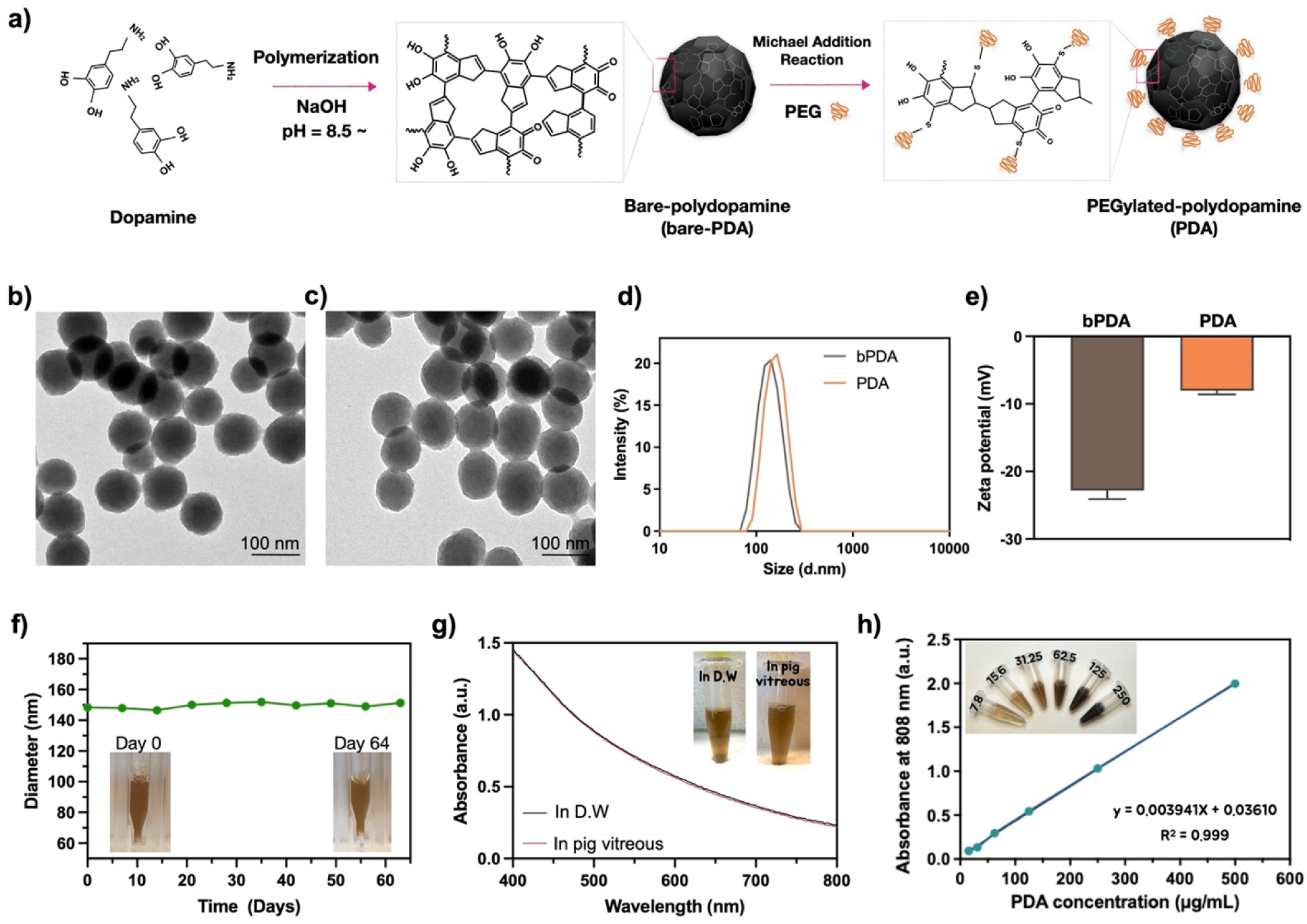
Characterization of polydopamine nanoparticles (PDA). a) A brief description of the polymerization and surface modification process of PDA. Transmission electron microscopy (TEM) images showed the morphology of purified b) bare-PDA and c) PDA (scale bar = 100 nm). d) The size distribution and e) zeta potential of bare-PDA and PDA were determined by DLS. Detection of the colloidal stability of PDA in pig vitreous without any f) aggregation and g) degradation for at least 9 weeks. The inset displays corresponding photographs of PDA dispersed in pig vitreous and aqueous solution. h) The absorbance at 808 nm for various concentrations of PDA showed a good linear relationship in the aqueous solution.

### 2.2 Evaluation of the scavenging activity of PDA

PDA has emerged as an antioxidant with a similar ROS-scavenging ability to natural RPE melanin[19,20,22]. Given that blue light induces the overproduction of ROS in retinal cells[23,24], the ROS-scavenging ability of PDA becomes especially important in suppressing ROS-induced cell damage. To assess the antioxidant capacity of PDA, we employed the DPPH and Evans Blue bleaching assay. The scavenging activity of free radicals was first assessed by measuring the absorbance change at 517 nm after mixing PDA with DPPH. As shown in **Figure 2a**, PDA exhibited effective antioxidative activities at the different concentrations (5 – 150 ug/mL), which aligned with the observed variations in solution colors following reaction of DPPH with different concentration of PDA **(Figure S4a supporting information).** A deeper yellow color in the solution indicated a lower remaining DPPH content, providing further evidence of the radical scavenging efficiency of the PDA. Similarly, the Evans Blue bleaching assay revealed the reduction of peroxynitrite levels in the presence of PDA by observing the fading of the blue color that indicates peroxynitrite-mediated oxidation **(Figure S4b supporting information)**. PDA effectively inhibited peroxynitrite levels by 39.8 % at a concentration of 150 μg mL−1 in comparison to the PBS control **(Figure 2b)**. Both assays demonstrated the oxidative activity of PDA as effective scavengers of ROS. We further analyzed the electron paramagnetic resonance (EPR) spin-trapping technique to quantify the catalytic production of short-lived, small free radicals. The magnetic parameters of the EPR spectra were detected after hydrogen peroxide was added to the reaction mixture containing DMPO spin trap, PDA, and iron sulfate to initiate the formation of hydroxyl radicals (OH^•^) in the course of the Fenton reaction. As shown in **Figure 2c**, the EPR signal decreases with an increasing concentration of PDA, becoming completely undetectable at 16 μg of PDA concentration, in comparison to the control group with no added PDA. Additionally, the plotted peak-to-peak amplitudes of EPR signals showed the time decay of the EPR intensity for the DMPO-OH^•^ spin adduct signal generated in the presence and absence of PDA, which clearly suggests that PDA is significantly effective as a radical scavenger **(Figure 2d)**. We conducted additional EPR spectroscopy to monitor whether PDA can exhibit the potential to function as a natural RPE melanin. The EPR signals clearly revealed that PDA is potential antioxidant with effective ROS scavenging activity compared with natural RPE melanin **(Figure S5 supporting information)**.

**Figure 2.**
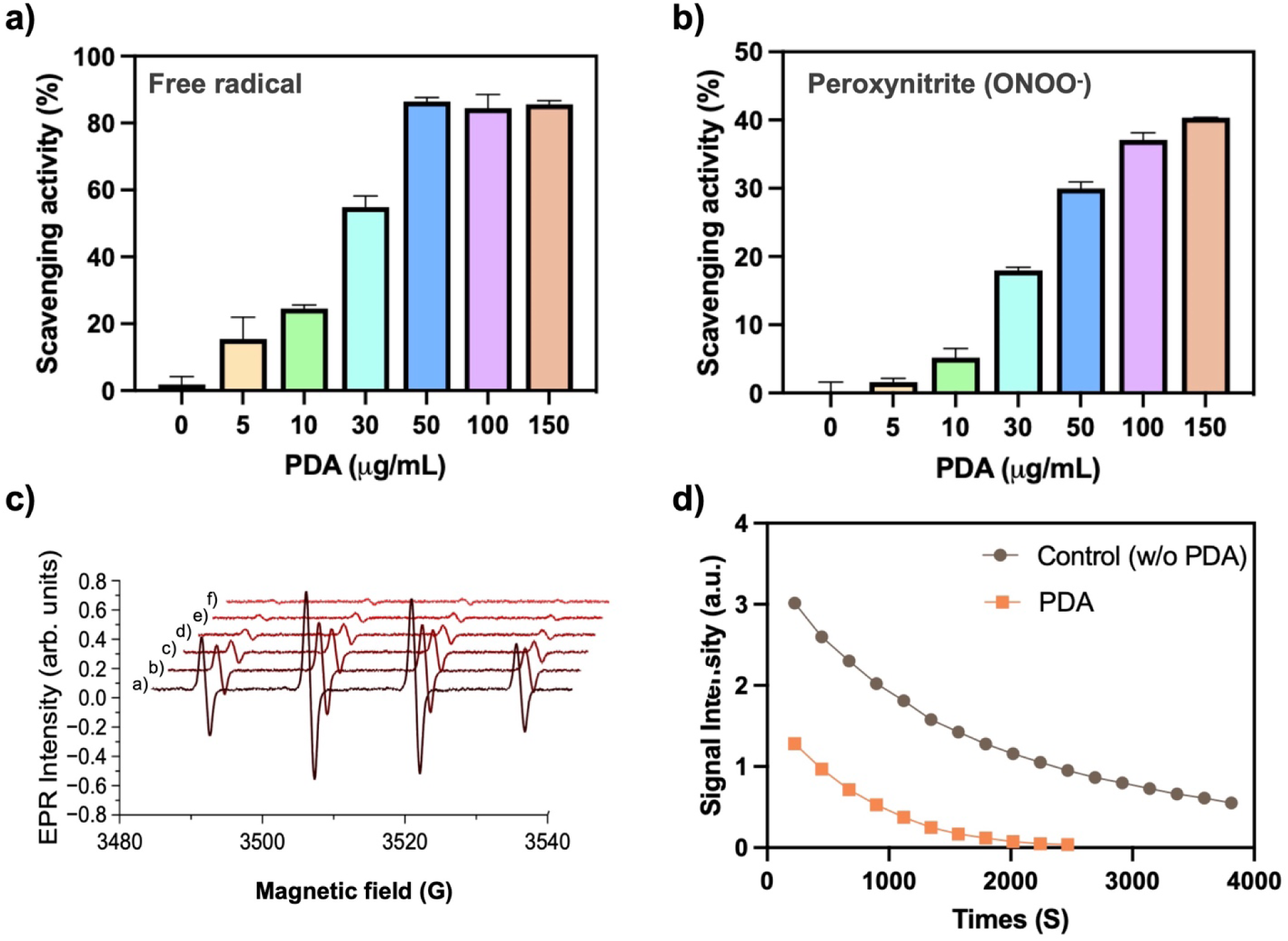
ROS-scavenging activity and EPR studies of PDA. a) Free radicals inhibition by PDA, and b) capacity of PDA to scavenge peroxynitrite radicals at different concentrations. c) Sequential EPR spectra of DMPO-OH• spin-adducts detected after a solution including 29.6 mM DMPO, 0.238 mM Fe(II) sulfate, and 10 mM H_2_O_2_ was treated with different amounts of PDA: (a) 0 μg mL^−1^ (no treatment); (b) 2 μg mL^−1^; (c) 4 μg mL^−1^; (d) 8 μg mL^−1^; (e) 16 μg mL^−1^; and (f) 32 μg mL^−1^. d) Peak-to-peak amplitude of EPR signal from DMPO-OH• adducts at 294 K in the absence (control, without PDA) and in the presence of 10 μg mL^−1^ PDA.

### 2.3 PDA-mediated protection on blue LEDs light-stimulated intracellular oxidative stress and phototoxicity

The broadband wavelength absorption capacity of melanin contributes to the protection of the retina from photodamage caused by ultraviolet (UV) and high-energy visible (HEV) light, including blue light[25,26]. To investigate the photoprotective effect of PDA against blue LEDs light-induced apoptosis in pigmented and non-pigmented pig RPE (pRPE) cells, we isolated pRPE cells from both normal and albino pig eyes **(Figure 3a)**. The pRPE cells were then exposed to blue light in a cell culture incubator with blue LEDs light system, as shown in **Figure 3a and Figure S6a supporting information**, and were assessed cell viability at various time points, including 1, 3, 6, and 24 hours of exposure to blue light. The blue light exposure exhibited time-dependent toxicity of pRPE cell growth, affecting both pigmented (brown bar) and non-pigmented (orange bar) pRPE cells. **(Figure 3b)**. The non-pigmented pRPE cells showed less than 60% survival at 6 h compared to pigmented pRPE cells, while the PDA-treated group (green bar) had a minimal impact on the survival of pRPE cells (80 ∼ %) under blue light exposure, which could be attributed to the photoprotective activity of PDA against blue light-induced apoptosis. Moreover, the bright-field images clearly supported that pigmented pRPE cells, when preincubated with PDA, exhibited a synergistic photoprotective effect by natural RPE melanin and PDA, enabling to maintain their morphology and survival in the exposure of blue light compared to other groups **(Figure 3c, Figure S7 supporting information)**. The suppression of ROS production by PDA was investigated by detecting intracellular ROS levels, which were measured using 2’,7’-dichlorodihydrofluorescein diacetate (DCFDA) that is known to produce green fluorescence in the presence of intracellular ROS[27]. The pRPE cells were first exposed under blue light exposure system for 6 h to induce oxidative stress by phototoxicity. Intracellular ROS levels exhibited a significant 3.5-fold increase in only blue light-stimulated pRPE cells (yellow bar) compared to DCFDA-only controls (purple bar, no blue light exposure), as shown in **Figure 3d**. Furthermore, the presence of PDA in pRPE cells resulted in a significant reduction of intracellular ROS production upon exposure to blue light (5 – 30 ug, *p <0.05, and ***p < 0.001). We conducted additional experiments to determine whether PDA can affect ROS stimulation in the absence of blue light exposure. The results revealed that the fluorescence levels were comparable to DCFDA (reagent only)-treated cells (**Figure S8 supporting information**). These results showed that the PDA does not induce any intracellular ROS.

**Figure 3.**
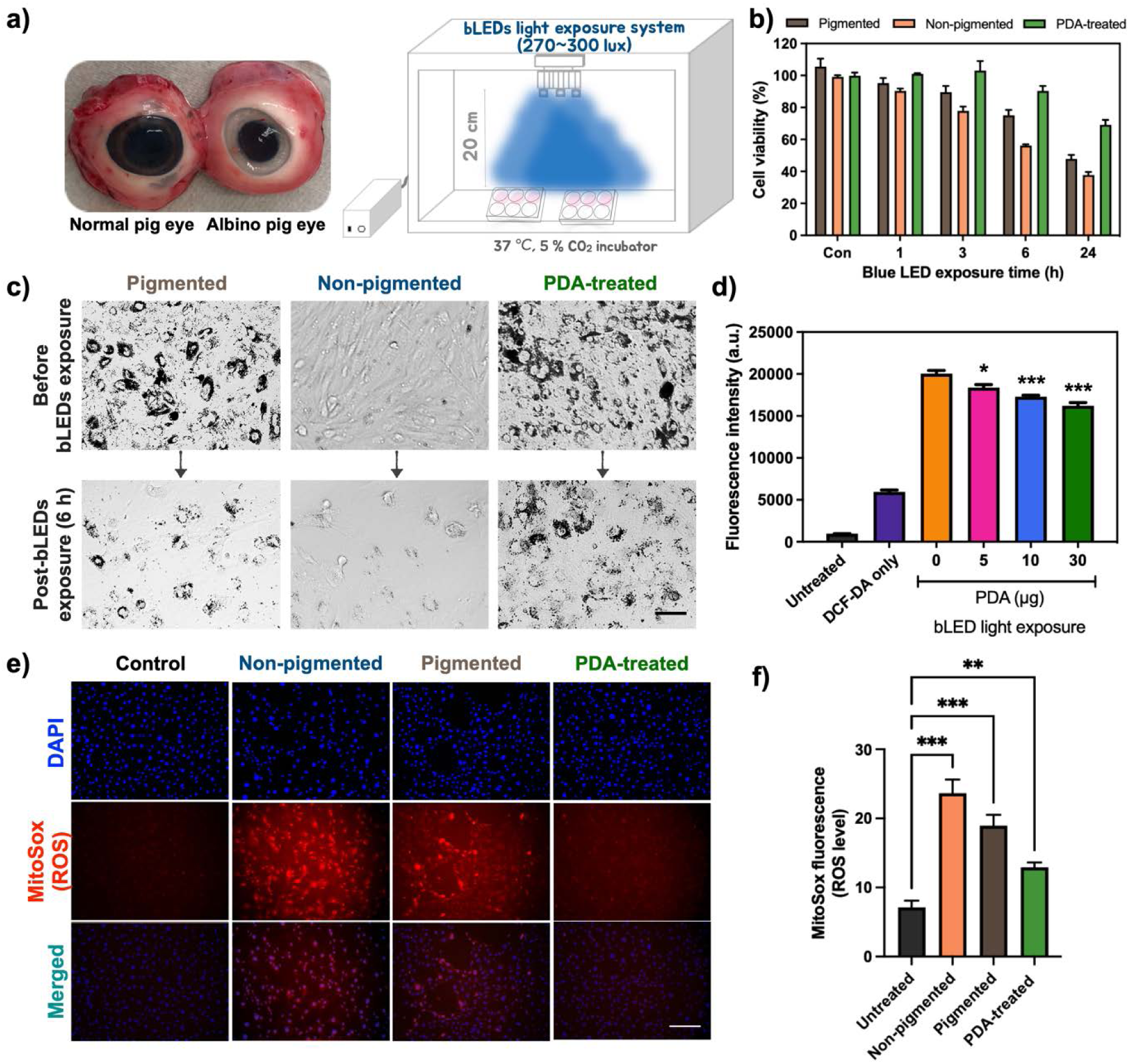
Photoprotective effect of PDA in primary pig RPE (pRPE) cells against a blue light-induced phototoxicity. a) The images of normal (pigmented) and albino (non-pigmented) pig eyes, and schematic illustration of blue LEDs light apparatus in the cell culture incubator for in vitro test. b) Cell viability (n = 3), following stimulation with blue light exposure in pigmented, non-pigmented, and PDA-treated pRPE cells. c) a significant decrease in the number of viable cells were found in non-treated pRPE cells, whereas PDA pretreatment significantly protected the blue light-exposed pRPE cells from the photocytotoxicity. d) Intracellular ROS scavenging activity of PDA in pRPE cells using DCFH-DA (50 μM). e) Representative MitoSox images exhibited blue light-stimulated oxidative stress in live pRPE cells in the absence and presence of PDA and showed that f) PDA treatment significantly (p*** < 0.001) reduced blue light-stimulated intracellular oxidative stress. Results presented from three independent experiments as mean ± SEM were analyzed with one-way ANOVA followed by Tukey’s post hoc multiple comparison test, *P < 0.1, **P < 0.01, ***P < 0.001, n = 3.

We next assessed the antioxidative activity of PDA using MitoSox Red staining, a fluorescent dye selective for mitochondrial-specific superoxide in live cells **(Figure 3e)**. We observed that the production of ROS was significantly higher in blue light-exposed non-pigmented pRPE cells (***P < 0.001) compared to pigmented pRPE cells. In contrast, the PDA-treated group showed a notable protective effect against oxidative stress induced by blue light phototoxicity **(Figure 3f)**. Hence, our results conclusively reveal that the presence of PDA in pRPE cells exhibits more effective photoprotective activity against blue light exposure than no treatment. Additionally, we were concerned about whether PDA can penetrate into the mitochondria and nucleus in the pRPE cells after reaching the cytosol. To address this safety issue, we conducted bio-TEM analysis to confirm that PDA does not interfere with the typical functionality of ROS during oxygen metabolism in the mitochondria. The bio-TEM images disclosed that internalized PDA exhibited a distinct localization within the cytoplasm, exhibiting no penetration into the mitochondria and nucleus in pRPE cells **(Figure S9 supporting information)** without any adversely impact cell viability **(Figure S10 supporting information)**.

### 2.4 Prevention of blue light-induced phototoxicity in Nrf2-deficient mouse model

Atrophic AMD phenotypes, such as drusen-like deposits, GA, inflammation, and retinal function impairment, are observed in Nrf2-deficient mice[28,29]. The downregulation of Nrf2 expression has the potential to compromise the effective regulation of oxidative stress, thereby rendering the retina more susceptible to blue light-induced oxidative stress. To investigate whether blue light exposure can lead to retinal atrophy, we selected pure Nrf2-deficient mice by genotype assay **(as shown in Figure S11 supporting information)** and exposure the mice to periodic blue light for 3 hours per day over 3 days (460 nm, 15,000 lux) using Kwon’s blue LEDs light exposure device, as described in the diagram in **Figure 4a and Figure S6b supporting information**. We first injected that of saline (1 μL/eye) and PDA (1 μL from a 1 μg/μL PDA stock) via a single intravitreal (IVT) administration, prior to inducing blue light-induced phototoxicity in Nrf2-deficient mice. Subsequently, we performed fundus fluorescein angiography (FFA) and optical coherence tomography (OCT) on day 5 and day 14 after blue light exposure to confirm photodamage in the retinal tissues. The representative images of FFA and OCT in the saline-injected group revealed that blue light exposure triggered photodamage in Nrf2-deficient mice, leading to permanent retinal impairment as retinal degeneration (**Figure 4b**). In contrast, the PDA-injected group was shown a condition closer to normal compared to that of the saline-injected group, indicating photoprotective ability of PDA against blue light-induced photodamage (**Figure 4c**).

**Figure 4.**
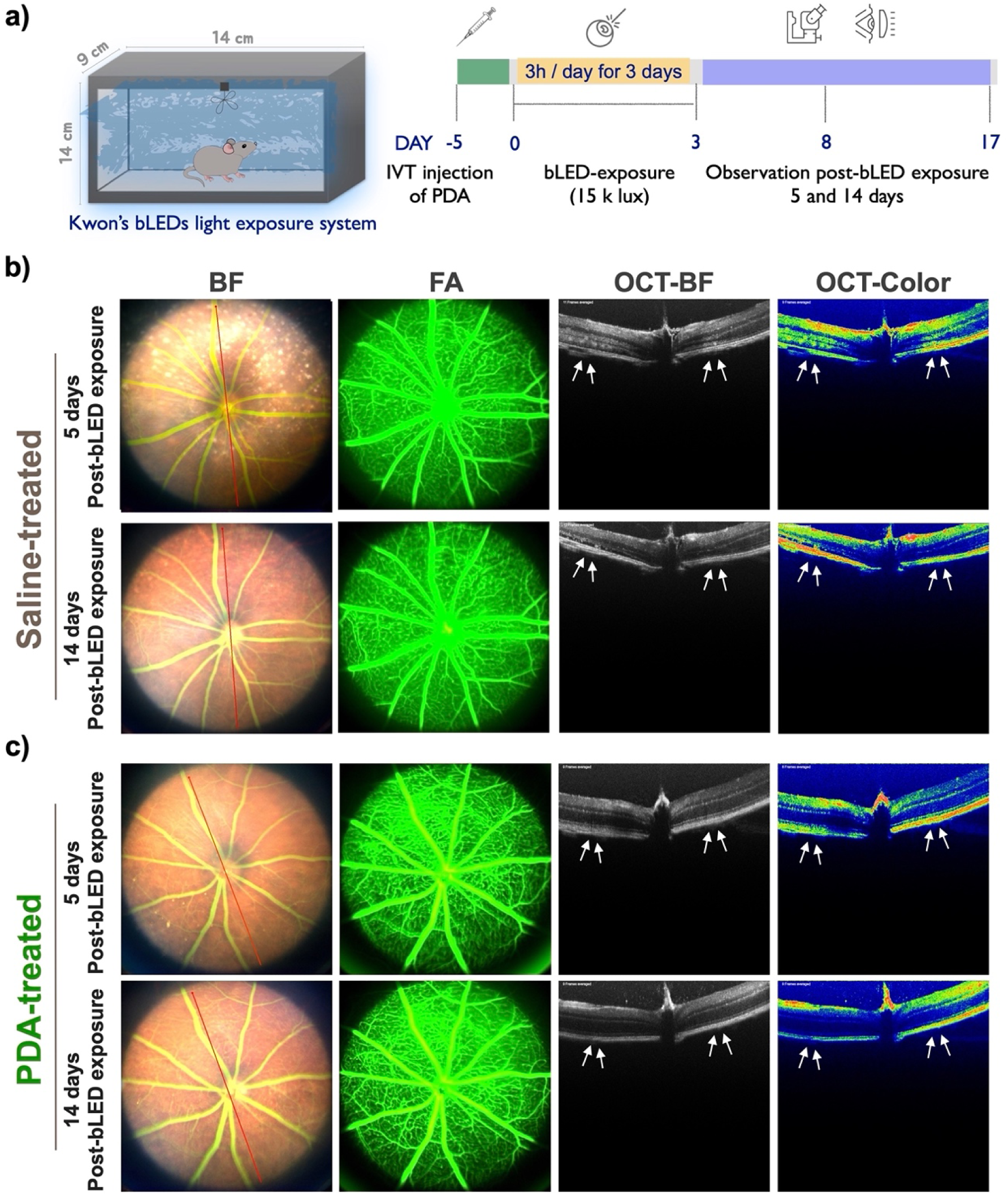
Fundus and OCT analysis of the effects of PDA administration in an Nrf2-deficient mouse model following blue light-induced phototoxicity. a) Schematic diagram of the periodic blue light (460 nm, 15K lx) exposure to Nrf2-deficient mice. 6 – 8 weeks Nrf2-deficient mice were exposed for 3 h per day for 3 days. At 5 days before blue light exposure, mice were intravitreally (IVT) injected with b) saline (1 μL) and c) PDA (1 μL from a 1 μg μL^−1^ PDA stock solution), respectively. Fundus and OCT photography were performed at day 5 and 14 post-injection. White arrows indicate areas of photodamage. Fluorescein angiography: FA, and bright field: BF.

### 2.5 The PDA protected retinal tissues from structural deformation by bule light-induced photodamage in Nrf2-deficient mice

We also assessed retinal histological analysis on Nrf2-deficient mice at 14 days after blue light exposure to bolster the FAA/OCT results and to quantitatively analyze the results. Histological studies revealed that blue light exposure in the saline-injected group led to severe damage and shrinkage in retinal tissue compared to the control group (no blue light exposure). However, the PDA-injected group showed no morphological changes induced by blue light-induced photodamage (**Figure 5a**). In addition, we evaluated the layer distribution of retinal tissues, including the inner plexiform layer (IPL), inner nuclear layer (INL), outer nuclear layer (ONL), and RPE layer, through statistical analysis of the thickness of each layer. The blue light exposure significantly shrank the thickness of the total retina in the saline-injected group, indicating irreversible photoreceptor cells degeneration. In contrast, the PDA-injected group did not cause any progress in the abnormality of retinal tissues, including the IPL (45.36 ± 1.77 μm), INL (32.76 ± 5.47 μm), ONL (48.65 ± 3.68 μm), and RPE (8.18 ± 0.11 μm), suggesting that PDA replenishment could prevent retinal degeneration against blue light-induced phototoxicity **(Figure 5b – f).**

**Figure 5.**
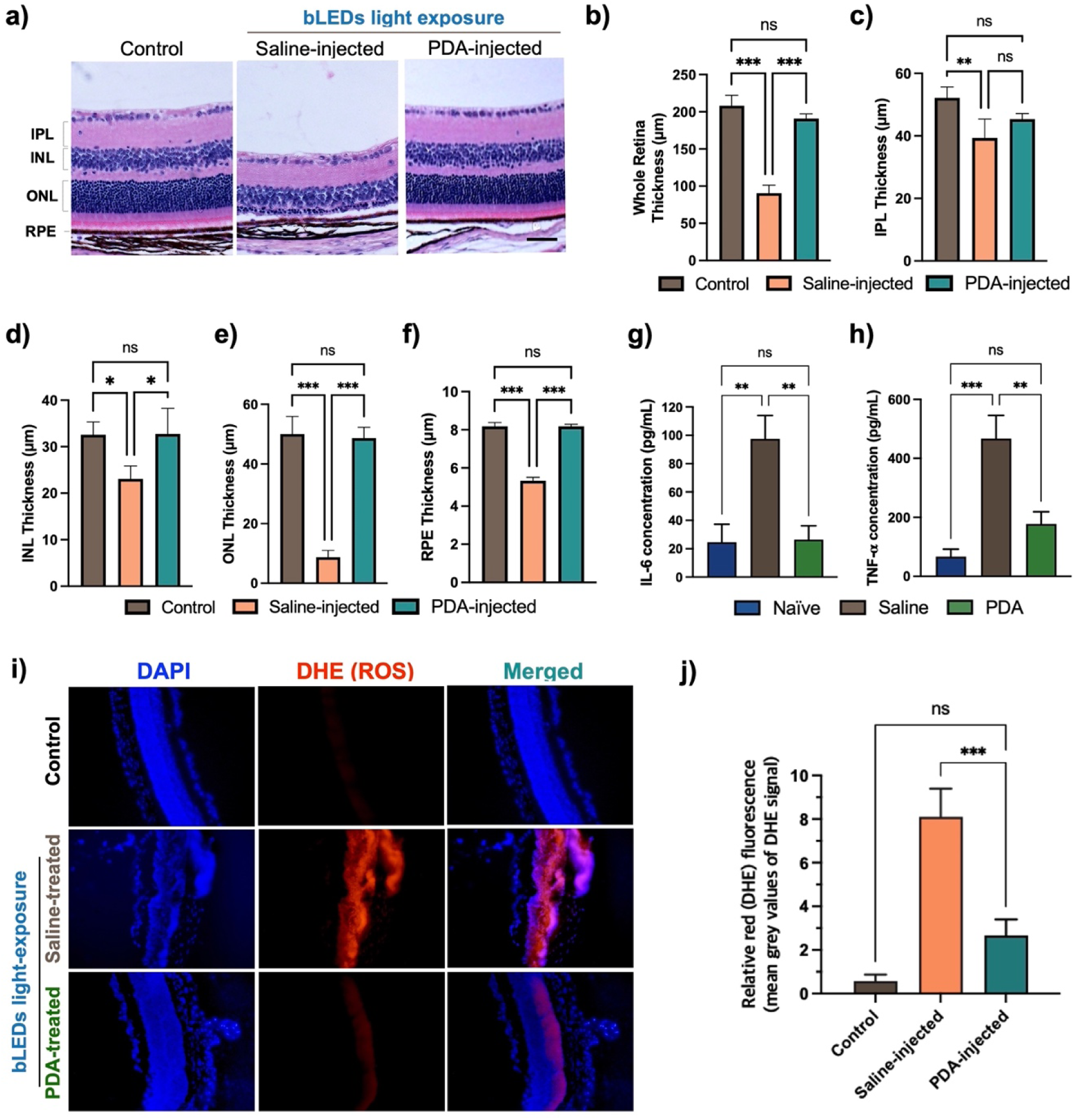
Evaluation of retinal tissues structure- and cytokines production-related markers, and detection of ROS levels in ocular tissues by dihydroethidium (DHE) staining post blue light exposure. a) H&E histology studies exhibited irreversible retinal damages by blue light exposure, and protective effect in the presence of PDA. Scale bar: 10 µm. Quantitative measurements were conducted to determine the thickness of b) the whole retina, c) IPL, d) INL, e) ONL, and f) RPE layers, serving as representative indicators of the retinal layer status. The expression levels of pro-inflammatory cytokines of g) tumor necrosis factor-α (TNF-α) and h) interleukin-6 (IL-6). i) Representative images of DHE staining in retinal tissues under blue light-induced phototoxicity, and j) the relative fluorescence intensities of ROS production. The values are the average of triplicate samples, and the data are represented as the mean ± SEM. Statistical significance was determined by a one-way ANOVA followed by Tukey’s post hoc multiple comparison test (* p < 0.05, ** p < 0.01, *** p < 0.001, ns, nonsignificant).

Retinal degeneration by phototoxicity is accompanied by an increased production of pro-inflammatory cytokines[30,31]. Therefore, we explored whether PDA plays a significant role in inhibiting relevant pro-inflammatory cytokines. We observed that blue light exposure in the saline-injected group induced overexpression of pro-inflammatory cytokines, specifically IL-6 **(Figure 5g)** and TNF-α **(Figure 5h)**, whereas replenishment with PDA significantly decreased the intensity of cytokine levels (***p<0.001), which was close to normal levels. Additionally, western blot analysis revealed that the presence of PDA significantly inhibited the expression levels of hypoxia-inducible factor (HIF-1α) and the oxidative marker protein (4-HNE adduct) in blue light-induced RPE/choroidal tissue compared to saline-injected group, as shown in **Figure S12a and b supporting information**.

A dihydroethidium (DHE) staining assay, extensively used to examine ROS production, was further performed to investigate blue light-induced ROS formation in ocular tissues. The blue light exposure stimulated high level of ROS production in the retinal layer, which was prominent compared to the PDA-injected group **(Figure 5i)**. The red intensity of ROS production was measured per tissue using ImageJ software. The PDA-injected group exhibited a significant reduction (***p<0.001) compared to the saline-injected group, suggesting that PDA suppresses ROS elevation and prevents photodamage **(Figure 5j)**.

### 2.6 Direct IVT delivery of PDA is safe and has no toxicity to ocular tissues

To evaluate potential ocular toxicity or morphological changes induced by PDA, we injected PDA into the vitreous space of Nrf2-deficient mouse eyes that had not been exposed to blue light. We measured the ONL thickness of the inferior and superior hemispheres at day 5 **(Figure 6a)** and day 60 **(Figure 6b)** post-IVT injection to monitor retinal pathological changes related to visual function and assess ocular tissue toxicity. No significant differences in saline and PDA injection were found between the groups. We also assessed intraocular pressure (IOP) changes at different time points, and no significant difference was observed **(Figure 6c)**. Additionally, electroretinogram (ERG) measurements were conducted at day 5 and 60 post-injection to assess the electrical activity of the retina. The ERG amplitudes of the PDA-injected group, specifically for a-waves in the range of 100 – 200 µV and b-waves in the range of 200 – 300 µV, showed no significant difference compared to the saline-injected group **(Figure 6d and e)**. Finally, we investigated the morphological changes by H&E staining at day 5, and 60 of post-injection to study the effect of PDA on morphology in the retinal cells. We observed no significant difference in the retinal nuclear cell density of the INL and ONL, as well as in the morphological changes, as shown in **Figure 6f, g, and h**. These results indicate that PDA are safe and have no toxicity to the eyes.

**Figure 6.**
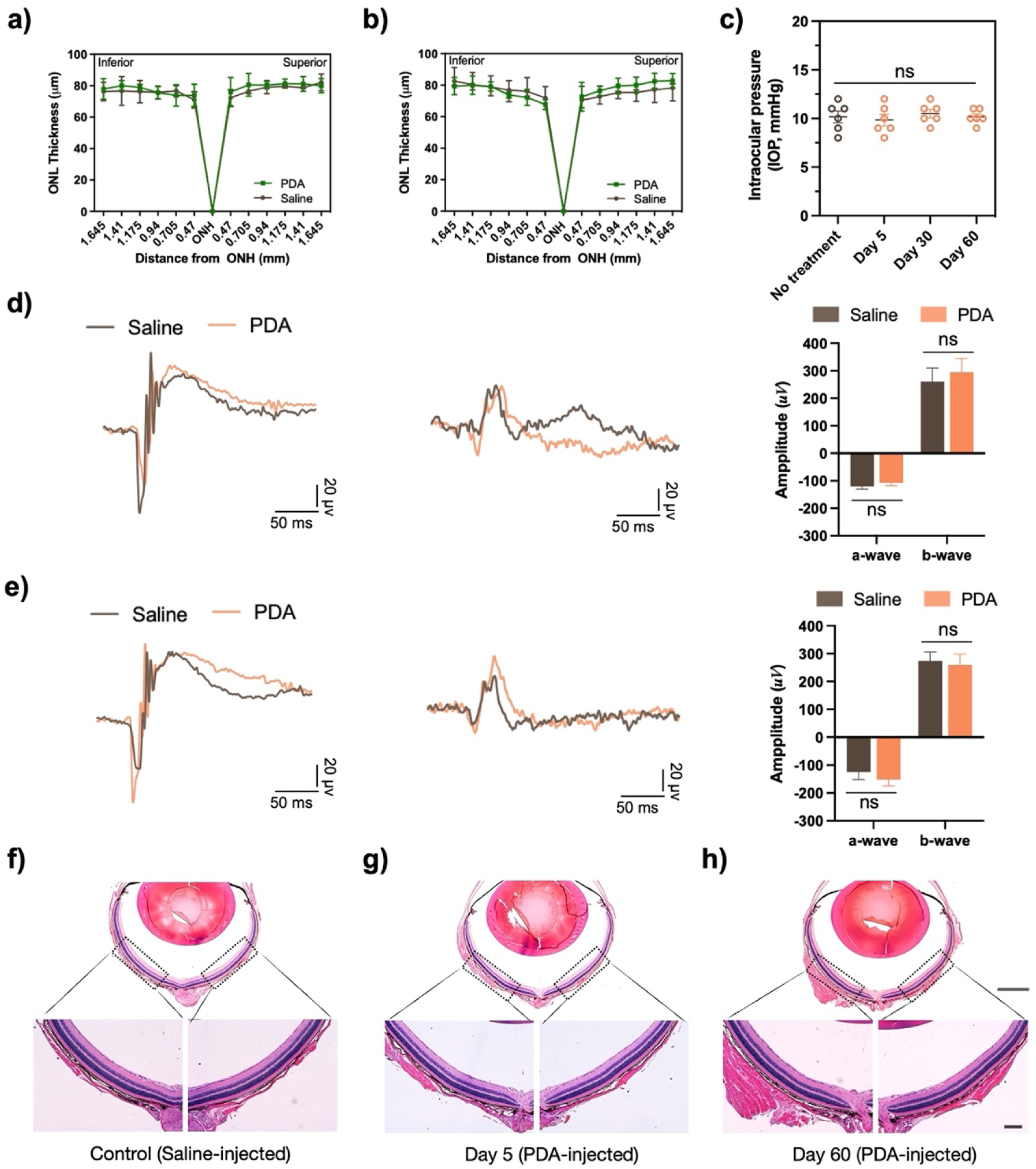
Safety evaluation of PDA in ocular tissues via IVT administration. Quantification of morphological changes in the outer nuclear layer (ONL) at a) day 5 and b) 60 of post-PDA injection, revealed no significant changes in ONL thickness between control (no injection), and PDA-injected eyes. c) Intraocular pressure (IOP) tested at day 5, 30 and 60 of post-PDA injection. Representative images of electroretinogram (ERG) test show dark-adapted and light-adapted waveforms from the saline- and PDA-injected groups at days 5 and 60 post-injection are shown in d) and e). f - h) Histological H&E analysis of retinas in (f) saline- and PDA-treated groups g) at day 5 and (h) at day 60 post-injection. Scale bar: 50 and 1000 µm as indicated. The data are represented as mean ± SEM, n = 5, and statistical analysis was performed using one-way ANOVA followed by Tukey’s post hoc multiple comparison test (n = 5, ns, nonsignificant).

## 3. Discussion

Blue light and its potential connection to ocular phototoxicity have been subjects of scientific investigation[4,32–34]. Ocular phototoxicity involves damage caused by exposure to light, particularly short-wavelength blue light, leading to harmful reactions in ocular tissues. Although ordinary ambient sunlight exposure does not cause phototoxicity, the concern is that prolonged or intense blue light exposure could contribute to ocular phototoxicity[32,35]. Short-wavelength blue light easily penetrates through the lens to reach the RPE cells, and then interacts with accumulated lipofuscin, as well as photosensitizers such as flavin and porphyrins within abundant mitochondria in retina cells[5,12]. Excessive exposure to blue light could result in the accumulation of ROS, adversely impacting RPE structure and function, ultimately leading to AMD disease.

The eyes possess inherent protective mechanisms against light-induced phototoxicity by the presence of melanin in ocular tissues[16]. In particular, RPE melanin plays a crucial role in the human eye as they protect the retina from oxidative stress and phototoxicity[15]. However, RPE melanin does not regenerate, and both the quality and quantity decrease with age, despite its crucial role in protecting retinal cells against oxidative stress caused by blue light-induced phototoxicity[36–38].

In this study, we present a rationally engineered and advanced strategy for replenishing melanin in the RPE. PDA, a synthetic polymer inspired by natural melanin, has emerged as an attractive therapeutic nanomaterial due to its photoprotection, biocompatibility, and antioxidant activities[18,20]. Moreover, the in vivo trafficking study via single-dose IVT injection demonstrated PDA prefer to accumulate in the RPE layer for up to 3 months, as found in an earlier study[19]. Therefore, we became more interested in seeking whether the replenishment of PDA in RPE cells could protect the eye against intense blue light exposure as an alternative and/or substitute for natural RPE melanin.

We prepared bare-PDA using a simple coordination and self-assembly strategy. Colloidal stability is critical to ensure that PDA remains dispersed in physiological fluids without forming aggregates, which could lead to undesired interactions with biological entities and compromise its overall performance[39]. Therefore, the surface of bare-PDA was successfully coated with PEG, reflecting a significant improvement in colloidal stability in pig vitreous for 60 days, as confirmed by DLS analysis **(Figure 1f)**. The damage to the RPE by accumulative oxidative stress is directly correlated with the pathogenesis of AMD. We observed that PDA effectively scavenged ROS, including free radicals, peroxynitrate, and hydroxyl radicals **(Figure 2)**. Furthermore, PDA demonstrated the ability to scavenge and/or prevent intracellular excess ROS production induced by blue phototoxicity in the pRPE cells **(Figure 3b, d, and e)**. Nrf2-deficient mice exhibit characteristics of atrophic AMD, but achieving the desired phenotypes in this knockout murine model takes a minimum of 12 months[40], presenting challenges for timely treatments. Studies suggest that combining multiple AMD risk factors in Nrf2-deficient mice may accelerate the development of an AMD-like model within a relatively short timeframe[29,41]. Our findings showed that blue light exposure contributed to the apoptosis and degeneration of retinal cells by triggering oxidative stress in an Nrf2-deficient mouse model **(Figure 4b, c and Figure 5a)**. Interestingly, PDA repressed the degeneration of retinal cells, consistent with earlier observations during in vitro photoprotection evaluations **(Figure 3c)**. Moreover, PDA-injected group demonstrated no significant signs of an inflammatory response against blue light-induced phototoxicity compared with saline-injected group **(Figure 5g and h)**.

The fact that PDA was explored as an alternative to natural RPE melanin without any associated toxicity to ocular tissues, is indicated by the ocular toxicity analyses in **Figure 6**. Altogether, our comprehensive studies demonstrated that replenishing PDA in RPE cells could protect retinal cells against oxidative stress caused by blue light-induced phototoxicity.

Given that ROS levels in AMD patients are higher than those in healthy individuals, the ROS scavenging ability and photoprotective effect of PDA could become especially important in suppressing ROS induced by blue light exposure. Our current study on the PDA replenishment strategy in the context of ocular phototoxicity by blue light exposure could contribute to an improved understanding of its functions and key role in related diseases, such as AMD.

## 4. Conclusion

In conclusion, we have demonstrated that PDA replenishment can serve as a natural antioxidant and a phototoxicity defense platform, protecting retinal tissues in the blue light-induced phototoxicity Nrf2-deficient mice model. Considering the role of oxidative stress in contributing to the pathogenesis of AMD, the replenishment of PDA presents a promising alternative approach in AMD prevention.

## 5. Material & methods

### 5.1 Synthesis and surface modification of PDA

Bare-PDA was synthesized using a previously published method with slight modifications[42]. Dopamine hydrochloride (180 mg) was dissolved in 90 mL deionized water, and 780 μL NaOH (1 N) solution was added under gentle stirring at 50 °C. Following a 5 h reaction, bare-PDA was purified through centrifugation at 13,000 rpm for 15 min. For surface modification, a 10 mL solution of bare-PDA (1 mg mL^−1^) was adjusted to pH 9 − 10 using NH_4_OH solution (28 wt%, Sigma-Aldrich). Subsequently, 50 mg of mPEG-SH (average Mn = 2000, Sigma-Aldrich) was introduced under continuous stirring for 12 h. After the reaction was completed, the PDA underwent a washing process through centrifugation at 13,000 rpm for 15 min with deionized water. Comprehensive characterization of MNPs, encompassing TEM, DLS, zeta potential, and UV-vis analysis, was conducted as outlined in the **Supporting Information section 1.2**.

### 5.2 Colloidal Stability

The colloidal stability of bare-PDA and PDA was assessed by preparing them through centrifugation (13,000 rpm, 20 °C, 15 min) and redispersing the pellet in pig vitreous. Measurements of colloidal stability were conducted using DLS analysis at weekly intervals.

### 5.3 Assessment of ROS scavenging efficiency of PDA

#### Free radical scavenging assay

The 2,2-diphenyl-1-picrylhydrazyl (DPPH•, Sigma–Aldrich) assay was performed following a previously described method with slight modifications[20]. Diluted DPPH• working solution (0.1 mM, in 95% ethanol) was added to various concentrations of PDA solution, resulting in final PDA concentrations of 0, 5, 10, 30, 50, 100 and 150 μg mL^−1^, respectively. The PDA solution without the DPPH• working solution served as the control, with the absorbance of PDA measured at 808 nm. Following a 20 min incubation in the dark condition, the DPPH• scavenging efficiency of PDA was determined by measuring the absorbance of each well at 516 nm. The calculation employed the equation:

DPPH•-scavenging effect (%) = [1 – (Ai – Aj)/Ac] X 100%.

where Ac is the absorbance of the DPPH• working solution, Aj is the absorbance of the PDA solution, and Ai is the absorbance of the PDA and DPPH• mixture. All measurements were performed in triplicate.

#### Peroxynitrite scavenging assay

The peroxynitrite (ONOO-) scavenging was measured by an Evans blue bleaching assay in the absence and presence of PDA. First, various concentrations of PDA solution (0, 5, 10, 20, 30, 50, and 100 μg mL^−1^) were added to a working solution containing DTPA, NaCl, KCl, and Evans blue in 50 mM PBS (pH 7.4) and then mixed with the ONOO-solution. After 30 min incubation at room temperature, the absorbance at 611 nm was measured to calculate the scavenging efficiency, with all determinations performed in triplicate.

### 5.4 EPR Spin-Trapping Experiments

For hydroxyl radicals (OH^•^) spin-trapping experiments, a working solution containing H_2_O (58 µL), DMPO (20 µL, 29.6 mM), FeSO_4_ (10 µL, 0.24 mM), and various concentration of PDA (2 µL, 2, 4, 8, 16 and 32 µg) underwent sonication for 15 minutes using a VWT Model 75D bath sonicator and thorough vortexing. Subsequently, hydrogen peroxide (H_2_O_2_, 10 µL, 10 mM) was added, and the mixture was incubated for 5 minutes to form a hydroxyl radical adduct. In a control experiment, deionized water was added to the reaction mixture instead of PDA immediately after H_2_O_2_ admixture. EPR spectra were recorded using a Bruker ELEXSYS E500 spectrometer (Bruker Biospin, Billerica, MA, USA) operating at approximately 9.867 GHz (X-band). The typical data acquisition parameters were set as follows: modulation amplitude, 0.5 G; microwave power, 2 mW; scan width, 100 G; sweep time, 40.96 s; time constant, 20.48 ms; conversion time, 40.00 ms. All EPR spectra were measured at 21 °C and Peak-to-peak amplitudes of EPR spectra were plotted vs. the time.

### 5.5 Pig primary RPE cells culture and isolation of RPE melanin

Primary pig RPE (pRPE) cells were obtained from the Ophthalmology animal facility at UNC-CH from 6 – 8-month-old healthy wild-type pigs, and the primary pRPE cell culture was conducted using the following procedure. Briefly, the extracted pig eye globes were rinsed with 70 % ethanol and washed with PBS solutions. The cornea, lens, and retina were removed, and the posterior eyecup was incubated with trypsin-EDTA (0.25%, Gibco) at 37 °C for 20 minutes. Subsequently, the pRPE cells were harvested by gently scraping off Bruch’s membrane. The collected mRPE cells were rinsed twice with DMEM/F-12 (1:1) (Gibco) containing 10% (vol/vol) fetal calf serum (Gibco) and incubated in a 35-mm culture dish at 37 °C under 95% humidity and 5% CO_2_. After two days of cell culture, residual unattached pRPE cells and debris were gently removed with a PBS buffer solution, and then the cell culture medium (DMEM/F-12 (1:1)) was changed every 2 – 3 days. To collect melanin from porcine RPE cells, we followed the isolation method with slight modifications as described earlier[43]. Briefly, collected pRPE cells were suspended in a hypotonic buffer containing protease inhibitor cocktail (MilliporeSigma™ Calbiochem™, 53913110VL). The pRPE cells underwent disruption with a Dounce tissue homogenizer. The whole cell lysate was then centrifuged at 3,000 g for 5 minutes and re-suspended in a buffer solution containing 50 mM Tris-HCl and 150 mM KCl (pH 7.4). Subsequently, the cell lysate was layered on top of a discontinuous OptiPrep® gradient (50 %, 35 %, 30 %, 20 %, and 15 %) and centrifuged at 135,000 g for 1 h at 4°C (Beckman L8-80M Ultracentrifuge, USA). The melanin pellet was recovered, purified with deionized water three times by centrifugation (Thermo Scientific Sorvall ST 16R centrifuge, USA) at 13,000 rpm for 15 minutes. The RPE melanin was finally redispersed in deionized water and stored at − 20 °C temperature.

### 5.6 Detection of ROS in blue light-stimulated pRPE cells

The intracellular ROS levels were measured using 2′,7′-dichlorodihydrofluorescein diacetate (DCFDA) and MitoSox staining following the manufacturer’s instructions. Briefly, pigmented and non-pigmented pRPE cells were seeded in 48-well plates at a density of 2 x 10^4^ cells per well (n = 4). The pRPE cells were allowed overnight at 37 °C in a 5 % CO_2_ atmosphere, and then the cells were exposed to 200 μL of fresh DMEM containing 30 μg of PDA (or PBS in the control experiments). After 24 h of incubation, the cells were washed with PBS and underwent blue LEDs exposure system (460 nm, LED MOD XLAMP BLUE STARBOARD, Digikey, USA) for 1, 3, 6, and 24 h. Subsequently, 200 μL of fresh medium containing 50 μM DCFH-DA solution was added to each well, while the untreated cells served as a control. To assess ROS levels, DCF fluorescence intensity was measured using a spectrophotometer (Molecular Devices Spectramax M5) with excitation and emission wavelengths of 485 and 530 nm, respectively. To confirm the intracellular ROS levels in live pRPE cells exposed to blue light-induced phototoxicity, 5 μM (final concentration) of a fluorogenic probe (MitoSOX, Invitrogen™) was added to each well after incubation with PDA. Then, the oxidized MitoSOX of intracellular oxidation (red fluorescence) was measured at excitation/emission: 510 nm/580 nm.

### 5.7 Animal

Nrf2-deficient mice were purchased from the Jackson Laboratory for bule light-induced phototoxicity studies. All experiments were conducted in accordance with the policies of the Institutional Animal Care and Use Committee (IACUC) at The University of North Carolina at Chapel Hill (UNC-CH) and were maintained following the Association for Research in Vision and Ophthalmology (ARVO) statements regarding the use of animals in ophthalmic and vision research. All mice in experiments were 6 − 8 weeks old postnatal. Ketamine (100 mg kg^−1^)/Xylazine (15 mg kg^−1^) was used for anesthesia in all cases where anesthesia was required. Euthanasia was performed using isoflurane exposure and decapitation.

### 5.8 Induction of retinal photodamage by blue light exposure

Five days before blue light exposure, we prepared a PDA solution in a 10 μL nanofil syringe with a 33 G blunt-end needle. After anesthetizing the Nrf2-deficient mice and penetrating under the limbus with a 27 G needle, PDA (1 μg μL^−1)^ was intravitreally injected using a micro syringe pump at a 20 μL min^−1^ infusion rate. The mice were placed on a warm bed until fully awake and kept in a temperature-controlled dark room. Before blue light exposure to Nrf2-defucuebt mice, the mice were dark-adapted for 16 hours. Next, the pupils of the mice were dilated with 1% tropicamide (Bausch & Lomb Inc., Tampa, FL, USA) under dim red light and acute retinal damage was induced by subjecting the mice to 15,000 lx of diffuse blue LEDs lights (Super Bright LEDs, 78 lm/W, 64 W, 5050 SMD LEDs) in Kwon’s blue light exposure system for 3 h for 3 days **(Figure S5b, supporting information)**. The mouse eyes were imaged using a Fundus camera and OCT on day 5 and 14 post-blue light exposure.

### 5.9 ERG, IOP, DHE Staining, and Histology Scan

ERG and IOP were recorded and analyzed as followed well-established methods published previously[19]. DHE staining assay (superoxide indicator, Invitrogen, D11347, USA) was employed to detect the ROS production in the retinal tissues. Briefly, mouse eyes were embedded in Tissue-Tek O.C.T. compound (Sakura Finetek, CA, USA) without fixation, and promptly frozen on crushed dry ice to preserve the ocular tissues. Subsequently, the specimens were cryo-sectioned at a thickness of 10 μm in groups of three, and the sections were stained with 5 μM DHE at 37 °C for 30 min. The samples were mounted with DAPI (Vectashield, CA, USA) after gently washing with PBS. The resulting images were captured using an Axiocam MR 5 camera on an Axio Observer.D1 inverted microscope (Carl Zeiss, Norway). For histologic analysis, the collected eyes underwent fixation with 4% paraformaldehyde, and were carried out Hematoxylin and Eosin (H&E) staining by the UNC Center for Gastrointestinal Biology and Disease Histology Core Facility.

### 5.10 Western blotting and pro-inflammatory ELISA

To determine protein levels, a western blot assay was performed. RPE/choroid tissues were homogenized in 0.5 mL RIPA lysis buffer (Thermo Fisher, Cat. 89900) and heated at 95 °C for 5 min after mixing with sample loading buffer (Bio-Rad), fresh 2-mercaptoethanol (99%, Sigma– Aldrich), and 0.1% sodium dodecyl sulfate (SDS, Sigma–Aldrich). Lysates containing 20 μg of protein were separated by 10% SDS-polyacrylamide gels and transferred to a polyvinylidene fluoride (PVDF, 0.45 μm, Immobilon) membrane. The membrane was blocked for nonspecific binding with 5% milk in 1x TBST (Tris-buffered saline, 0.1% Tween 20) and incubated with rabbit polyclonal anti-4-HNE (ab46545, Abcam, 1:1500) and HIF-1α (141795, Cell Signaling, 1:1,000) in 5 % milk at 4 °C overnight. Subsequently, the membrane was washed 1x TBST (3 times, 10 min each) and incubated with goat anti-rabbit and anti-mouse IgG antibodies (sc-2030 and sc-2031, respectively, Santa Cruz) at 1:20,000 for 1.5 h at room temperature. The electrochemiluminescent signal was analyzed using a ChemiDoc™ MP imaging system (Bio–Rad), and densitometric analyses were carried out with Image Lab software v4.1 (Bio–Rad). Pixel densities in each band were normalized to the corresponding amount of beta-actin (Proteintech, Cat. 60004–1, 1:50,000) monoclonal antibody. The anti-inflammatory effects of PDA were assessed by quantifying the production of inflammatory cytokines, TNF-α and IL-6, utilizing commercially available ELISA kits, specifically the TNF-α ELISA kit (R&D Systems Inc., Minneapolis, MN, USA) and the IL-6 ELISA kit (R&D Systems Inc., Minneapolis, MN, USA), following the manufacturer’s instructions.

### 5.11 Statistical analysis

The results from all investigations are expressed as the mean ± standard deviation (S.D.) derived from a minimum of three independent experiments. Statistical and kinetic analyses were conducted using GraphPad Prism 9.0 software (La Jolla, CA, USA). Multiple group comparisons were assessed through one- and two-way ANOVA as deemed appropriate, with statistical significance established at a p-value below 0.05 (P < 0.05).

## Acknowledgements

The authors thank Ahra Cho (Biotechnology, Durham Technical Community College) for critical reading of the manuscript. The authors thank Amar Shankar Kumbhar (The Chapel Hill Analytical and Nanofabrication Laboratory, CHANL, UNC-CH) and Carolyn Batchelor Suitt (Histology Core, UNC Center for Gastrointestinal Biology and Disease Histology Core Facility) for kind help with the transmission electron microscopy imaging and H&E staining of ocular tissues. This work was supported by the Edward N. & Della L. Thome Memorial Foundation (138298, Z.H.) and the BrightFocus Foundation (M2022001F to Y.K.)

## References

[1] W. L. Wong, X. Su, X. Li, C. M. G. Cheung, R. Klein, C.-Y. Cheng, T. Y. Wong, Lancet Glob Health 2014, 2, e106.

[2] S. Jabbehdari, J. T. Handa, Surv Ophthalmol 2021, 66, 423.

[3] Y. Ozawa, Redox Biol 2020, 37, 101779.

[4] P. N. Youssef, N. Sheibani, D. M. Albert, Eye 2011, 25, 1.

[5] Z.-C. Zhao, Y. Zhou, G. T. and J. Li, Int J Ophthalmol n.d., 11, 1999.

[6] G. Tosini, I. Ferguson, K. Tsubota, Mol Vis 2016, 22, 61.

[7] C. J. Kennedy, P. E. Rakoczy, I. J. Constable, Eye 1995, 9, 763.

[8] J. R. Sparrow, M. Boulton, Exp Eye Res 2005, 80, 595.

[9] Z. Ablonczy, D. Higbee, D. M. Anderson, M. Dahrouj, A. C. Grey, D. Gutierrez, Y. Koutalos, K. L. Schey, A. Hanneken, R. K. Crouch, Invest Ophthalmol Vis Sci 2013, 54, 5535.

[10] R. A. Radu, N. L. Mata, A. Bagla, G. H. Travis, Proceedings of the National Academy of Sciences 2004, 101, 5928.

[11] L. E. Lamb, J. D. Simon, Photochem Photobiol 2004, 79, 127.

[12] J.-X. Tao, W.-C. Zhou, X.-G. Zhu, Oxid Med Cell Longev 2019, 2019, 6435364.

[13] J. Stone, D. van Driel, K. Valter, S. Rees, J. Provis, Brain Res 2008, 1189, 58.

[14] C. Núñez-Álvarez, C. Suárez-Barrio, S. del Olmo Aguado, N. N. Osborne, Acta Ophthalmol 2019, 97, e103.

[15] D.-N. Hu, J. D. Simon, T. Sarna, Photochem Photobiol 2008, 84, 639.

[16] M. Istrate, B. Vlaicu, M. Poenaru, M. Hasbei-Popa, M. C. Salavat, D. A. Iliescu, Rom J Ophthalmol 2020, 64, 100.

[17] T. Sarna, J. M. Burke, W. Korytowski, M. Różanowska, C. M. B. Skumatz, A. Zaręba, M. Zaręba, Exp Eye Res 2003, 76, 89.

[18] Y. Huang, Y. Li, Z. Hu, X. Yue, M. T. Proeto, Y. Jones, N. C. Gianneschi, ACS Cent Sci 2017, 3, 564.

[19] Y.-S. Kwon, M. Zheng, A. Y. Zhang, Z. Han, ACS Nano 2022, 16, 19412.

[20] Y.-S. Kwon, M. A. Voinov, M. Zheng, A. I. Smirnov, Z. Han, Nano Today 2023, 50, 101879.

[21] I. Singh, G. Dhawan, S. Gupta, P. Kumar, Front Microbiol 2021, 11.

[22] X. Lou, Y. Hu, H. Zhang, J. Liu, Y. Zhao, J Nanobiotechnology 2021, 19, 436.

[23] M. Abdouh, M. Lu, Y. Chen, A. Goyeneche, J. V. Burnier, M. N. Burnier, Exp Eye Res 2022, 217, 108978.

[24] P. Geiger, M. Barben, C. Grimm, M. Samardzija, Cell Death Dis 2015, 6, e1985.

[25] B.-L. L. Seagle, E. M. Gasyna, W. F. Mieler, J. R. Norris, Proceedings of the National Academy of Sciences 2006, 103, 16644.

[26] M. Xiao, M. D. Shawkey, A. Dhinojwala, Adv Opt Mater 2020, 8, 2000932.

[27] E. Eruslanov, S. Kusmartsev, in Advanced Protocols in Oxidative Stress II (Ed.: D. Armstrong), Humana Press, Totowa, NJ, 2010, pp. 57–72.

[28] Z. Zhao, Y. Chen, J. Wang, P. Sternberg, M. L. Freeman, H. E. Grossniklaus, J. Cai, PLoS One 2011, 6, e19456.

[29] K. Wang, M. Zheng, K. L. Lester, Z. Han, Sci Rep 2019, 9, 14573.

[30] G. Kaur, N. K. Singh, Neural Regen Res 2023, 18.

[31] N. Fernando, Y. Wooff, R. Aggio-Bruce, J. A. Chu-Tan, H. Jiao, C. Dietrich, M. Rutar, M. Rooke, D. Menon, J. T. Eells, K. Valter, P. G. Board, J. Provis, R. Natoli, Invest Ophthalmol Vis Sci 2018, 59, 4362.

[32] A. Cougnard-Gregoire, B. M. J. Merle, T. Aslam, J. M. Seddon, I. Aknin, C. C. W. Klaver, G. Garhöfer, A. G. Layana, A. M. Minnella, R. Silva, C. Delcourt, Ophthalmol Ther 2023, 12, 755.

[33] N. A. Wong, H. Bahmani, Heliyon 2022, 8, e10282.

[34] X. Ouyang, J. Yang, Z. Hong, Y. Wu, Y. Xie, G. Wang, Biomedicine & Pharmacotherapy 2020, 130, 110577.

[35] C.-H. Lin, M.-R. Wu, C.-H. Li, H.-W. Cheng, S.-H. Huang, C.-H. Tsai, F.-L. Lin, J.-D. Ho, J.-J. Kang, G. Hsiao, Y.-W. Cheng, Toxicological Sciences 2017, 157, 196.

[36] T. Sarna, J. M. Burke, W. Korytowski, M. Różanowska, C. M. B. Skumatz, A. Zaręba, M. Zaręba, Exp Eye Res 2003, 76, 89.

[37] T. Sarna, J Photochem Photobiol B 1992, 12, 215.

[38] U. Schraermeyer, J. Kopitz, S. Peters, S. Henke-Fahle, P. Blitgen-Heinecke, D. Kokkinou, T. Schwarz, K.-U. Bartz-Schmidt, Exp Eye Res 2006, 83, 315.

[39] J. Schubert, M. Chanana, Curr Med Chem 2018, 25, 4553.

[40] Z. Zhao, Y. Chen, J. Wang, P. Sternberg, M. L. Freeman, H. E. Grossniklaus, J. Cai, PLoS One 2011, 6, e19456.

[41] H. Huang, A. Lennikov, Exp Eye Res 2020, 196, 108061.

[42] K.-Y. Ju, Y. Lee, S. Lee, S. B. Park, J.-K. Lee, Biomacromolecules 2011, 12, 625.

[43] L. Pelkonen, M. Reinisalo, E. Morin-Picardat, H. Kidron, A. Urtti, PLoS One 2016, 11, e0160352.

