## Supplemental Information for "Polydopamine Nanoparticles as Mimicking RPE Melanin for the Protection of Retinal Cells Against Blue Light-Induced Phototoxicity"

**1. Experimental Section**

**1.1 Materials.**

Dopamine hydrochloride (Sigma-Aldrich, H8502), Sodium hydroxide (NaOH, Sigma-Aldruch, S8045), Poly(ethylene glycol) methyl ether thiol (mPEG-SH, Sigma-Aldrich, 729140), 2,2-Dipheny-1-picrylhydrazyl (DPPH, Sigma-Aldrich, D9132), peroxynitrite (Cayman, 81565), Evans Blue (Sigma-Aldrich, E2129), MitoSOX (Thermo Fisher Scientific InvitrogenTM, USA), Polybed 812 epoxy resin (Polysciences, Inc.). FeSO_4_ 7H_2_O was purchased from Fluka Chemie Gmbh (Buchs, Switzerland), 1-hydroxy-2-oxo-3-(N-methyl-3-aminopropyl)-3-methyl-1-triazene (NOC 7) and 5,5-dimethyl-1-pyrroline N-oxide (DMPO) were purchased from Sigma-Aldrich. Hydrogen peroxide was purchased from VWR (Radnor, PA). All chemicals were used as received without any further purification. Deionized water was used in all experiments.

**1.2 Material Characterization**

High-resolution scanning/transmission electron microscopy (S/TEM, Thermo Scientific^TM^ Talos F200X) was utilized for examining the morphology of the samples at the Chapel Hill Analytical and Nanofabrication Laboratory (CHANL, UNC-CH). TEM images were captured after the samples were collected on 200-mesh carbon-coated copper TEM grids. Additionally, the size distribution and zeta potential of the samples were analyzed by dynamic light scattering (DLS) using a Nano-ZS Zeta Sizer system (Malvern, USA) to complement TEM data and provide insights into the size changes of the PDA nanoparticles. Ultraviolet-visible light (UV-Vis) spectra were recorded on an Thermo Scientific™ NanoDrop™ OneC Microvolume spectrophotometer.

**1.3 Fundus fluorescein angiography (FFA) and optical coherence tomography (OCT) imaging procedure**

To assess morphological changes of retinal tissues following blue light exposure, FFA and OCT imaging procedures were conducted. The fluorescein angiography was performed according to established methods^[1,2]^. Briefly, mice were anesthetized through intraperitoneal administration of ketamine (85 mg kg^−1^) and xylazine (10 mg kg^−1^) mixture. Subsequently, intraperitoneal injection of 1 % AK-FLUOR (Alcon, 100 cc 20 g^−1^ mouse) was carried out for fluorescein angiography. Pupils were dilated using topical 1 % tropicamide (Bausch & Lomb Inc., Tampa, FL, USA), and mice were positioned on the Micron III fundoscopy system platform (Phoenix Research Laboratories, Pleasanton, CA, USA). Adjustments were made until clear images of FFA and OCT were visualized. The area of interest was captured using StreamPix software.

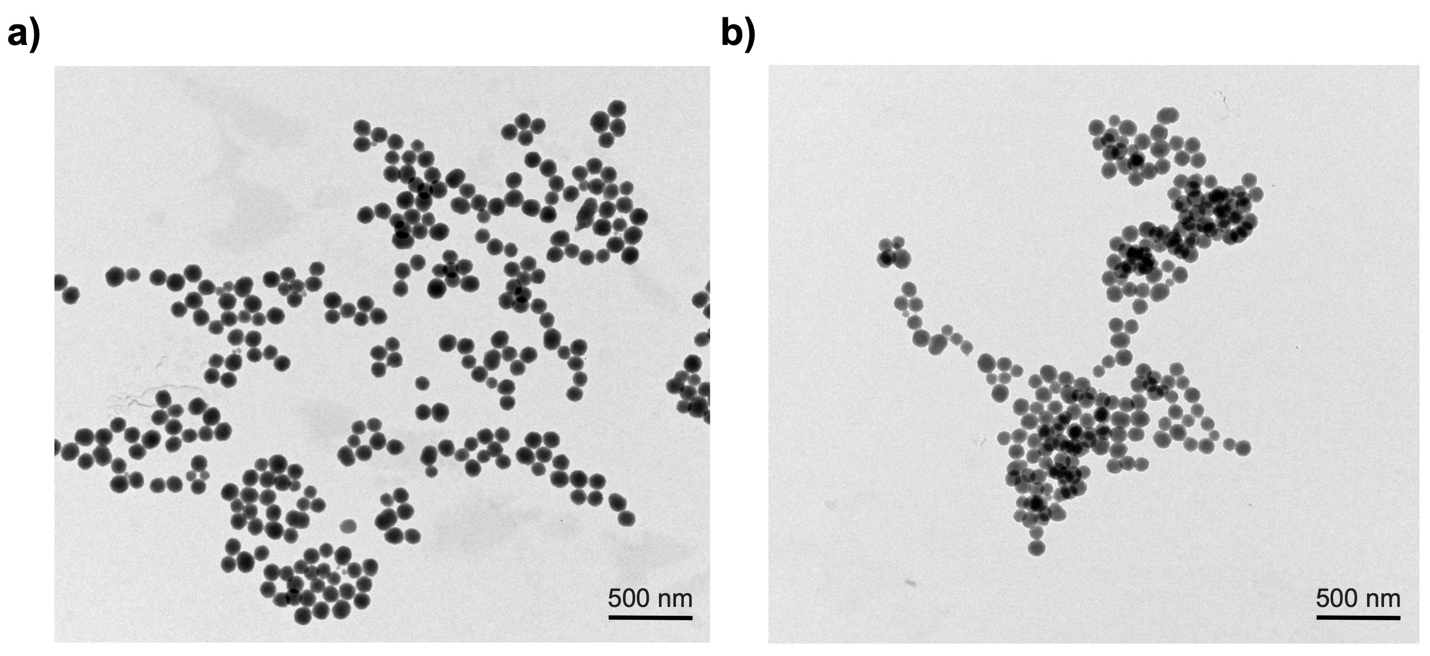

**Figure S1.** TEM images of a) bare-PDA and b) PDA exhibit a uniform shape and size at low magnification. No significant morphological differences in PDA were observed after surface modification with mPEG-SH.

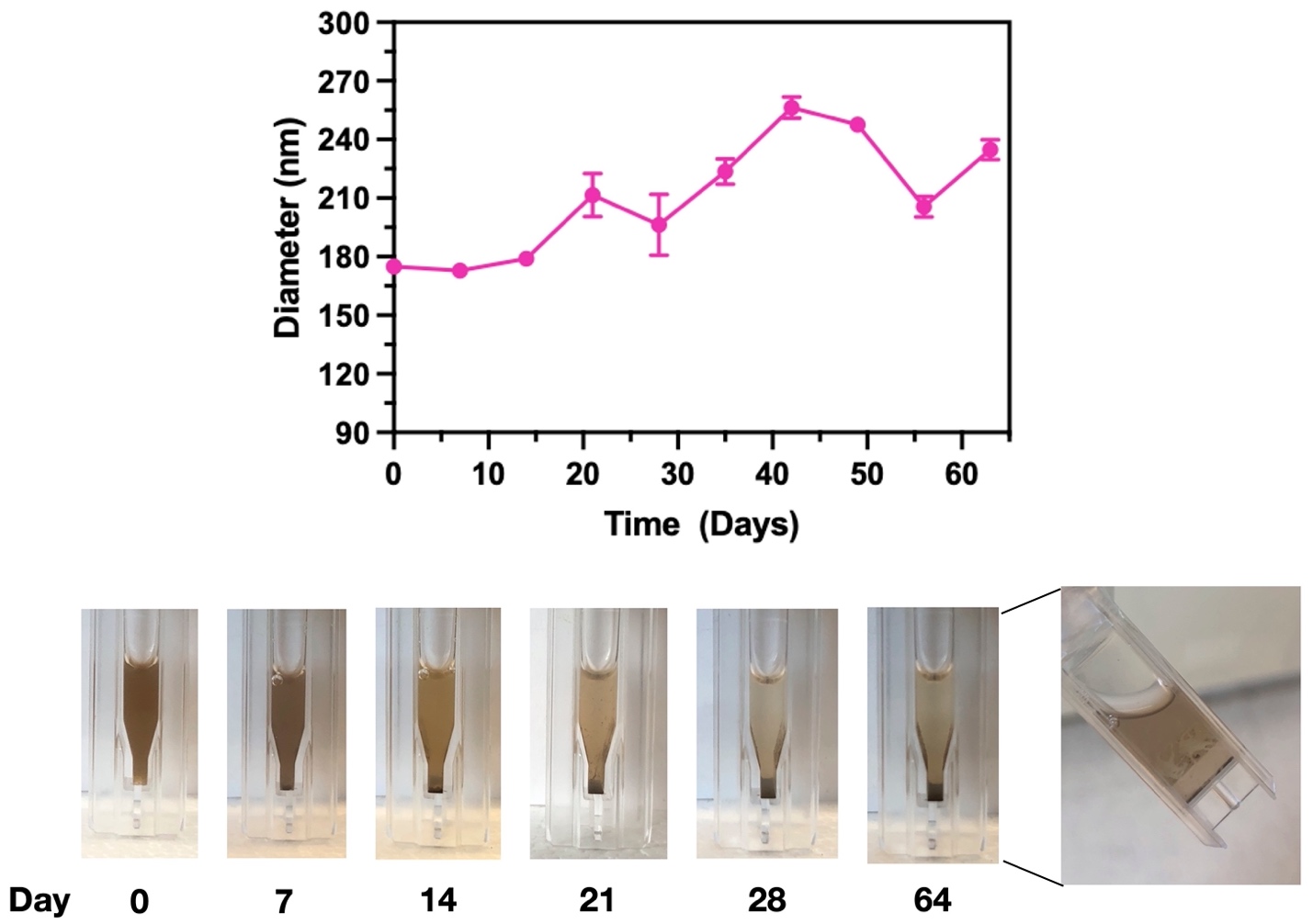

**Figure S2.** The in vitro colloidal stability test demonstrated that bare-PDA exhibited poor colloidal stability at various time points (top panel), which was clearly revealed in photographs of bare-PDA dispersed in pig vitreous (bottom panel).

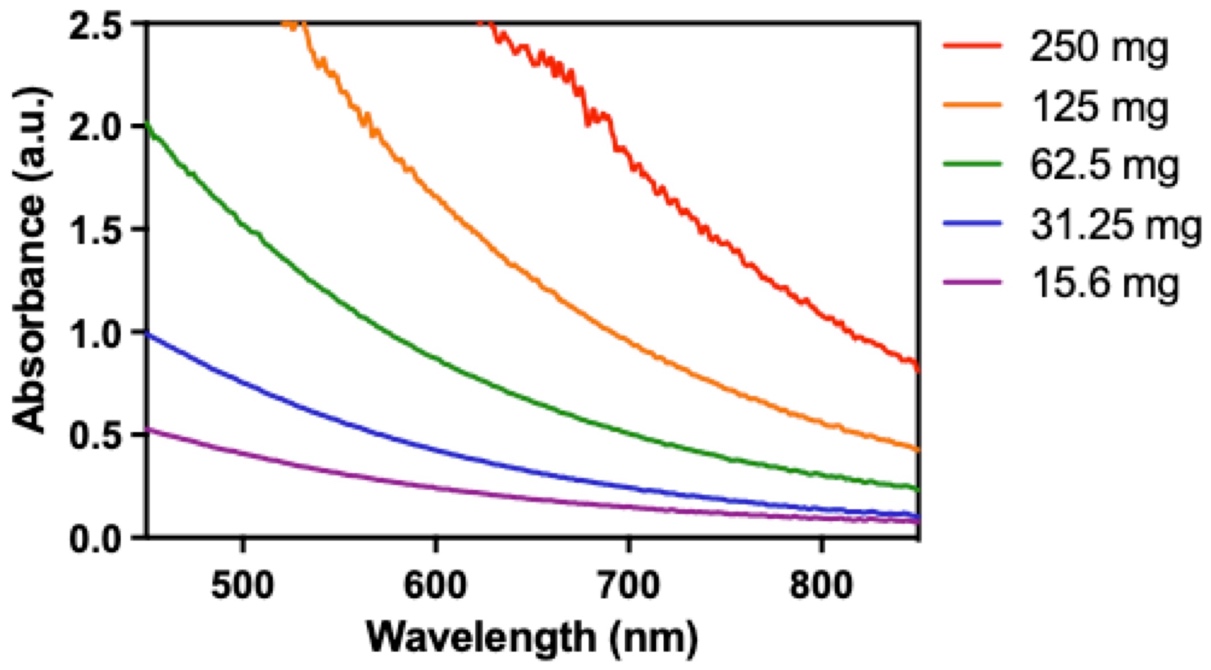

**Figure S3.** UV-vis absorbance spectra were obtained in PDA dispersions at various concentrations.

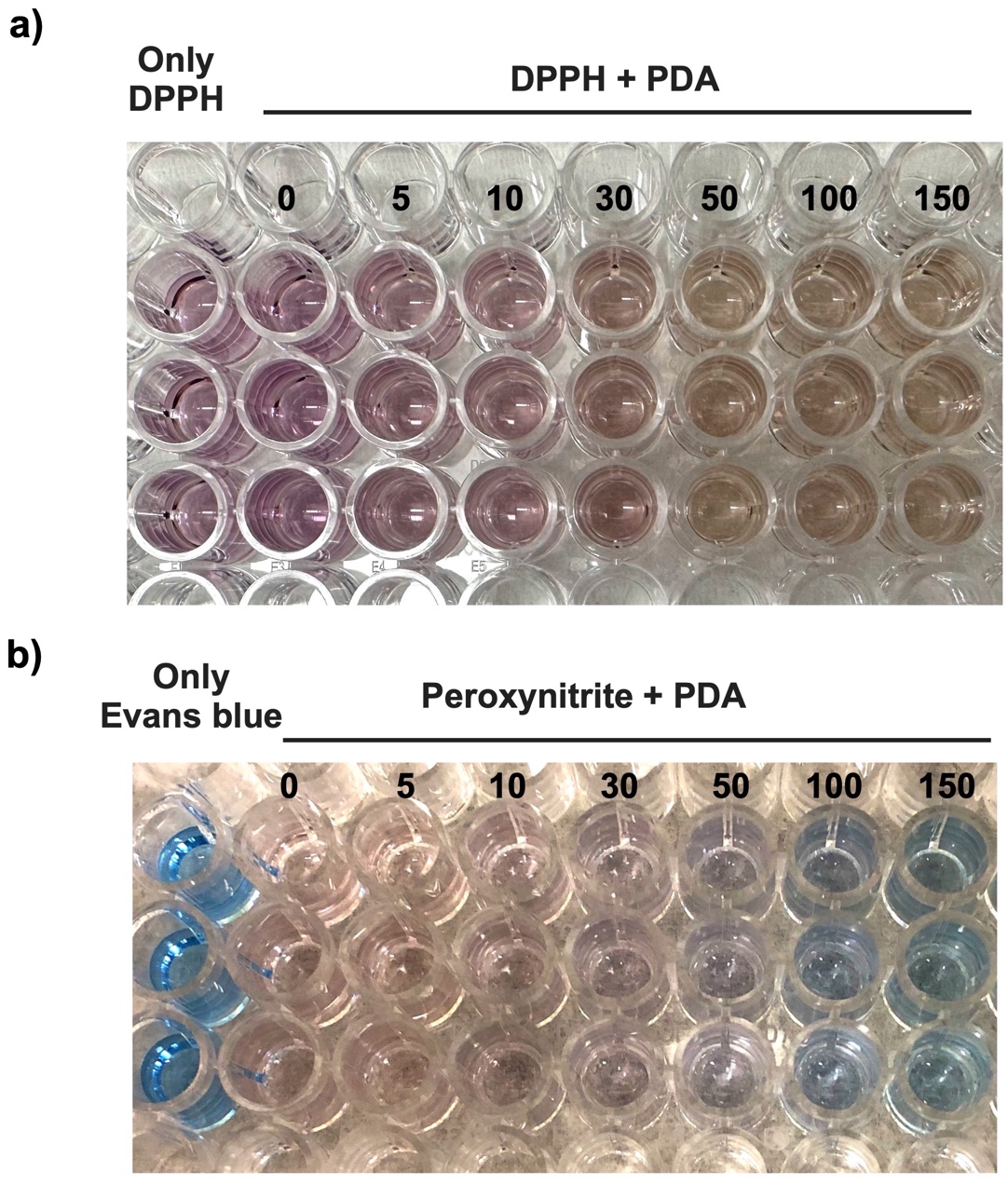

**Figure S4.** Representative photographs of DPPH and Evans blue assays demonstrated proportionate antioxidant capacity to different concentrations of PDA. The compound (DPPH^•^), a stable radical cation with a purple color, and Evans blue with peroxynitrite caused discoloration of the solution in accordance with the antioxidant activity of PDA.

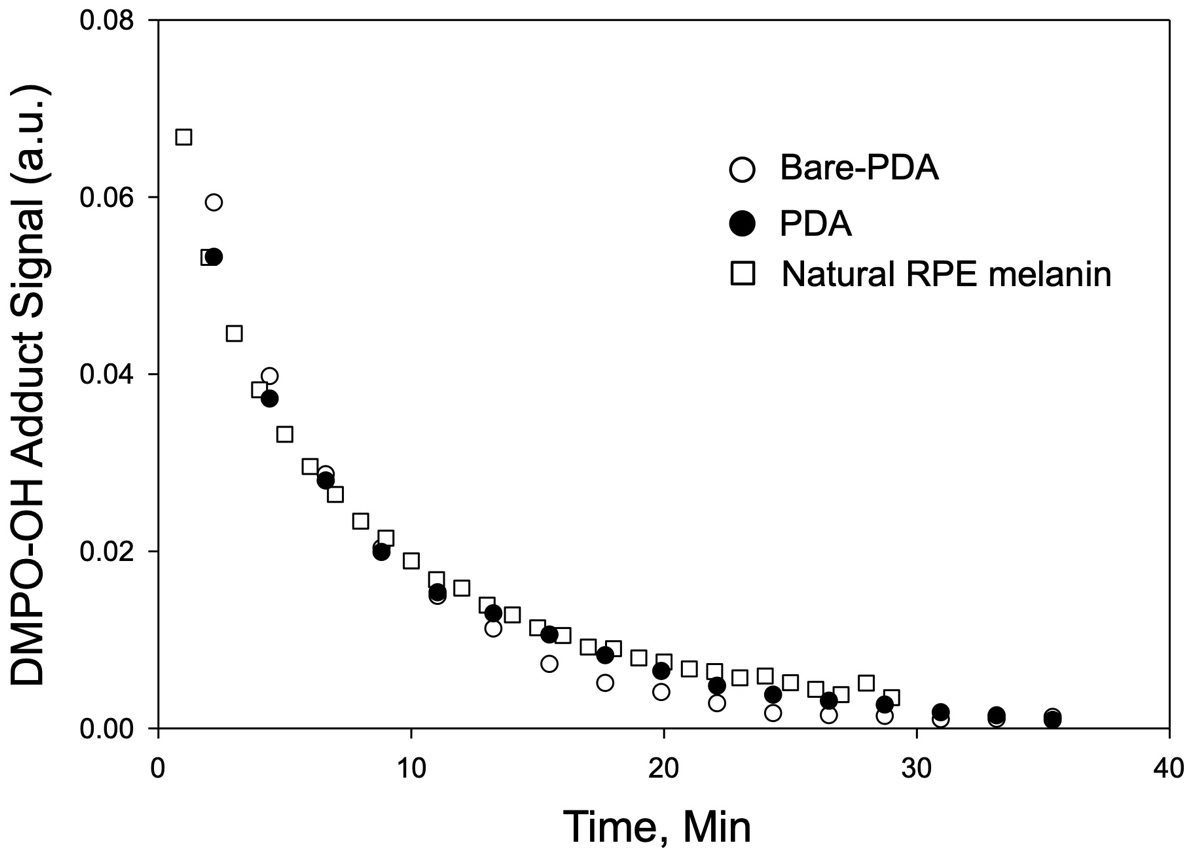

**Figure S5.** Peak-to-peak amplitude of the EPR signal from DMPO-OH• adducts at 294 K in the presence of bare-PDA (circles), PDA (filled circles) and natural RPE melanin (squares).

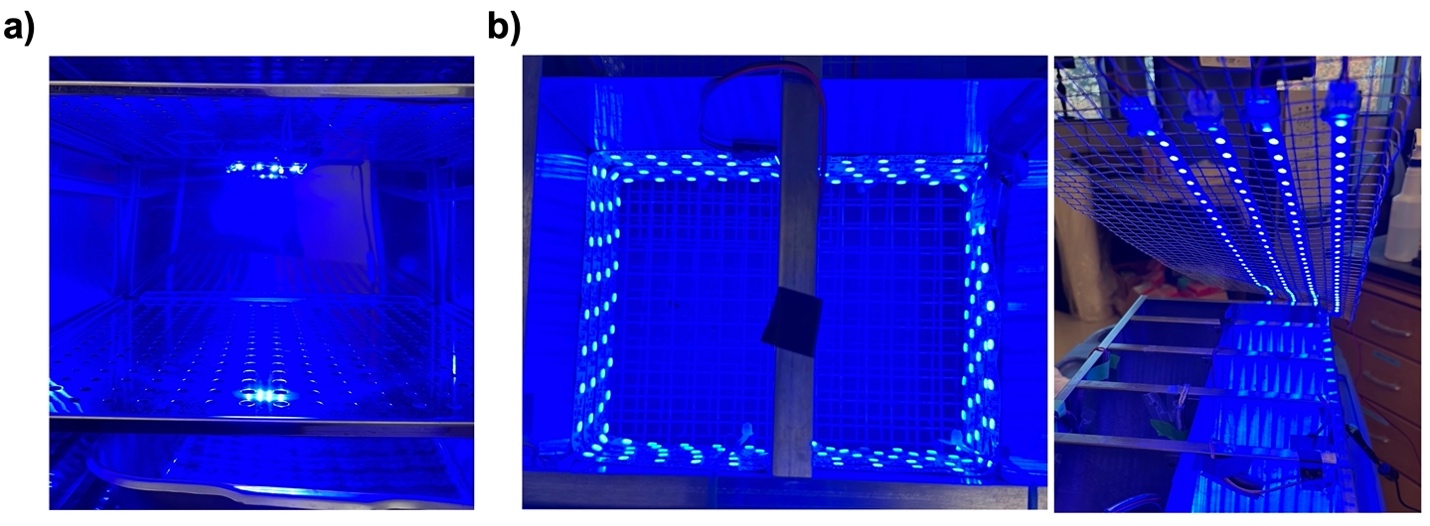

**Figure S6.** Representative images of Kwon's blue light exposure system. a) The intense blue LEDs light apparatus (460 nm, LED MOD XLAMP BLUE STARBOARD, Digikey, USA) were positioned on the top rack inside the cell culture CO_2_ incubator for in vitro experiments. b) The illuminated cages with blue LEDs light apparatus (Super Bright LEDs, 78 lm/W, 64 W, 5050 SMD LEDs) for in vivo test. The mice were not able to observe each other or interact and were maintained under controlled conditions (22 ± 1 °C, 60 ± 10 % relative humidity, with food and water provided ad libitum).

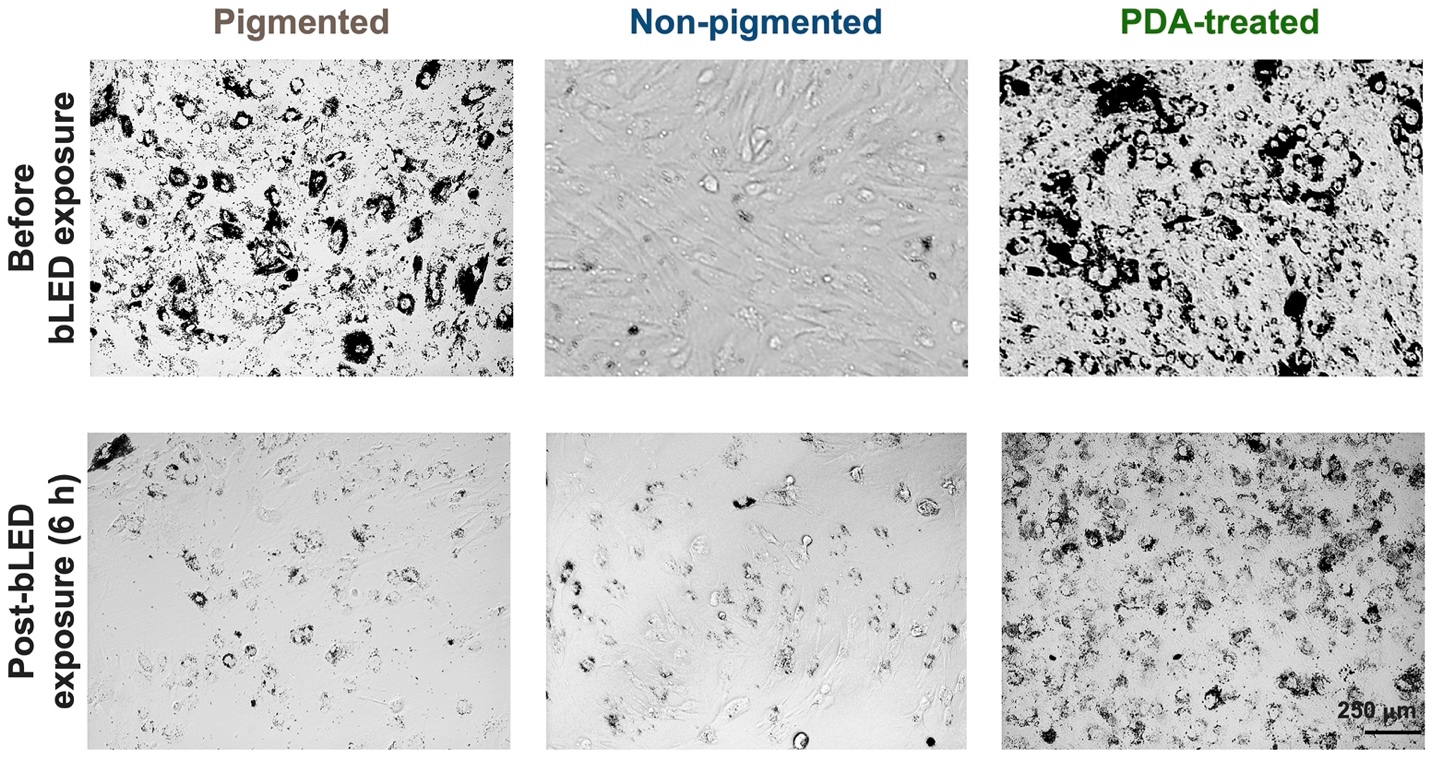

**Figure S7.** Representative bright-field images demonstrate the photoprotective activity of PDA against pRPE cells exposed to blue light during a 6-hour incubation, captured at low magnification. Scale bar: 250 μm.

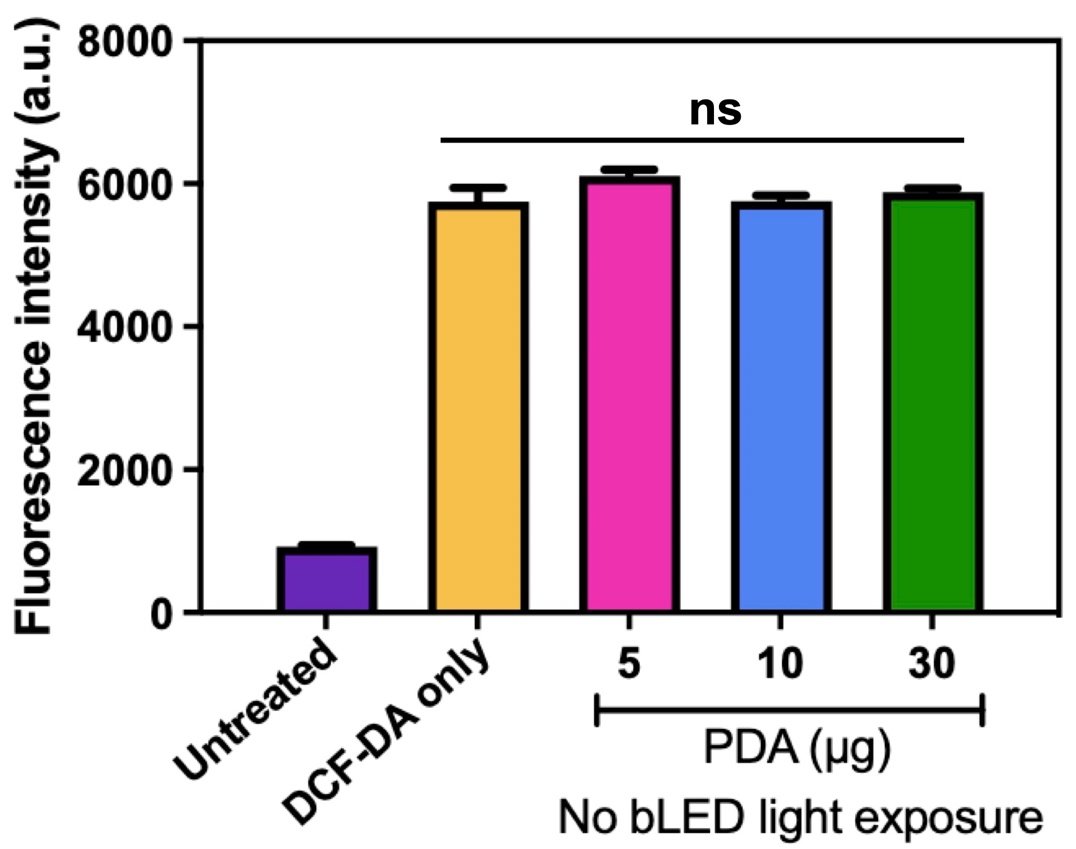

**Figure S8.** The intracellular ROS levels was determined in a separate experiment without blue light exposure. we observed no interactions between DCFDA and PDA (5 – 30 µg), indicating that PDA did not induce intracellular ROS production. ns, nonsignificant.

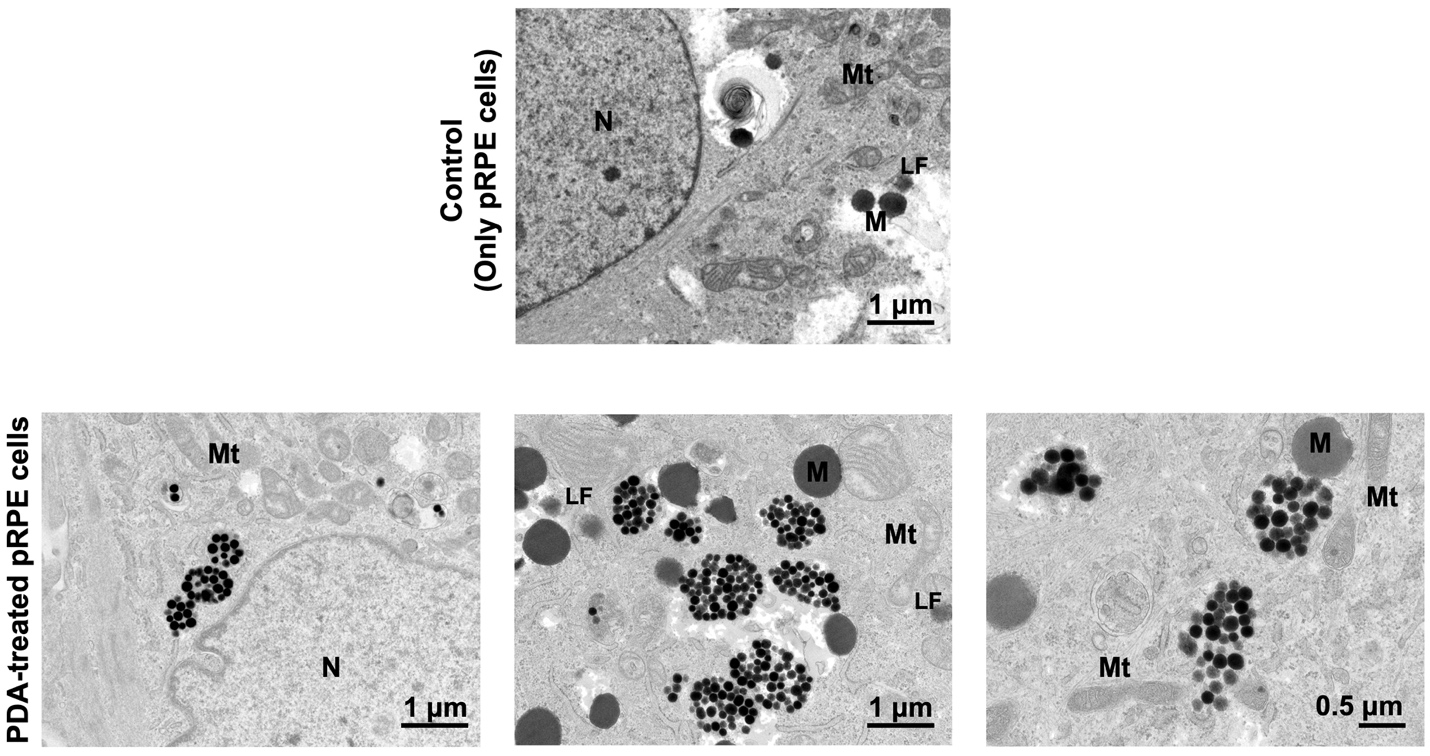

**Figure S9.** Bright-field transmission electron microscopy images of pRPE cells incubated with PDA showed that PDA is exclusively present in the cytosol of pRPE cells, with no detection in the nucleus and mitochondria (N; nuclear, Mt; mitochondria, M; melanin, and LF; lipofuscin).

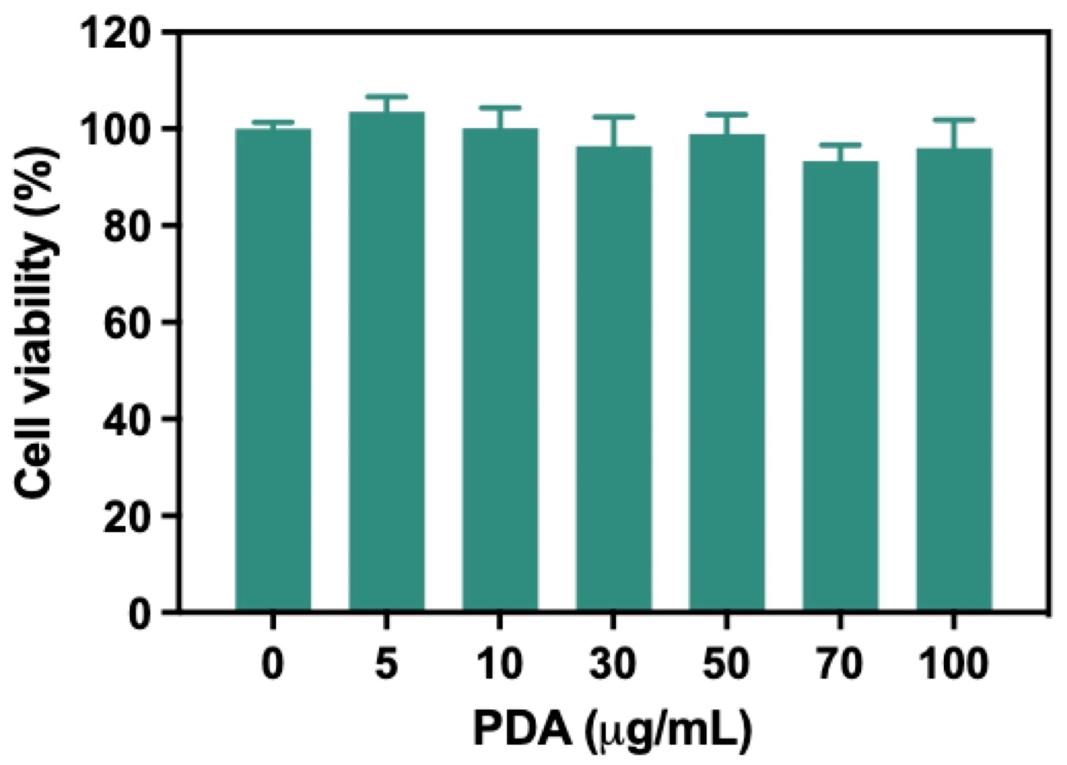

**Figure S10.** Cell viability test for PDA in pRPE cells demonstrated high cell viability (> 90 %) even after treatment with different concentrations (0 to 100 µg mL^−1^) for 24 hours. Results presented from three independent experiments as mean ± SEM were analyzed.

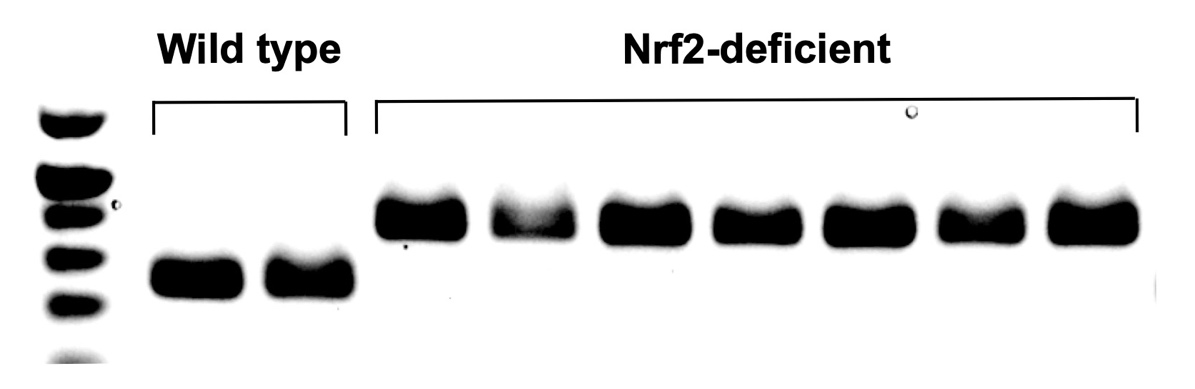

**Figure S11.** Genotype result was obtained from wild type and Nrf2-deficient mice. Pure Nrf2-deficient mice were selected to induce oxidative stress and phototoxicity through blue light exposure in this experiment.

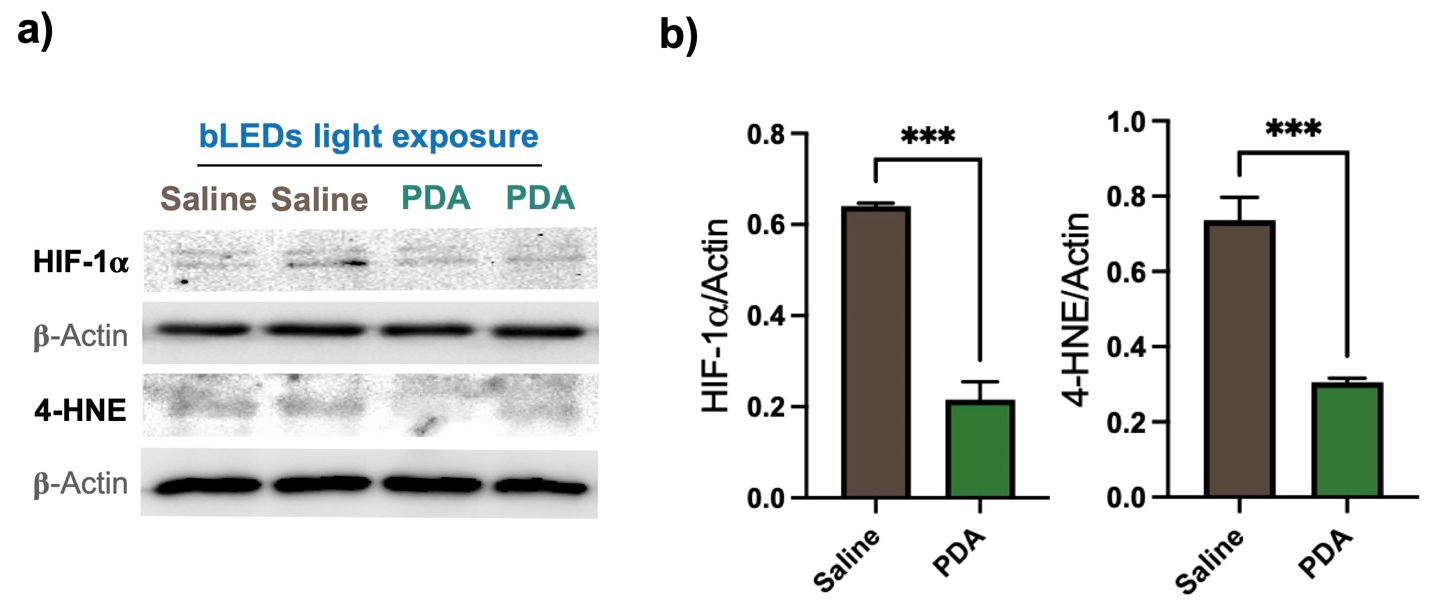

**Figure S12.** Western blot analysis was performed in RPE/choroid/scleral tissues lysates collected from saline- (as control), and PDA-injected groups after blue light-induced phototoxicity. a) Representative images of western blot analysis determining the expression levels of HIF-1a and 4-HNE. b) The relative expression levels of HIF-1a and 4-HNE were significantly reduced following treatment with PDA (***P < 0.001).

**References**

[1]	Y.-S. Kwon, M. Zheng, A. Y. Zhang, Z. Han, *ACS Nano* **2022**, *16*, 19412.

[2]	Y.-S. Kwon, M. A. Voinov, M. Zheng, A. I. Smirnov, Z. Han, *Nano Today* **2023**, *50*, 101879.
